# Elevated procoagulant platelets driven by necroptosis and pyroptosis aggravate pulmonary thrombosis via suppressing monocyte efferocytosis in severe pneumonia

**DOI:** 10.64898/2026.09.14.751620

**Authors:** Shi-Qing Li, Li-Yin Liao, Wei-Yang Fan, Jun-Bin Liang, Xi-Xian Liu, Meng-Yi Li, Chang-Yuan Kang, Hong-Xuan Zhou, Li-Chao Yang, Jin-Hua He, Zheng-Shi Lin, Xiao-Hong Chen, Shu-Feng Ma, Zhen-Hui Zhang, Zi-Feng Yang

## Abstract

**BACKGROUND:** Severe influenza pneumonia with secondary bacterial infection is complicated by progressive pulmonary thrombus exacerbation, a key contributor to respiratory failure, yet anticoagulant therapies show limited efficacy and bleeding risks. Although platelet-monocyte crosstalk initiates thrombosis, whether and how it drives thrombus exacerbation via procoagulant platelets and monocyte efferocytosis remains unclear.

**METHODS:** Clinical samples, mouse models, and isolated platelets challenged with influenza A virus followed by methicillin-resistant *Staphylococcus aureus* (MRSA) were analyzed. Procoagulant platelet, platelet programmed cell death and monocyte efferocytosis were assessed, and pharmacological inhibition, platelet depletion and systemic/platelet-specific *Gsdmd* knockout were used. Platelet proteomics and exogenous C1qa supplementation identified C1qa as a key mediator.

**RESULTS:** We showed that elevated procoagulant platelet-driven thrombus exacerbation, rather than initial thrombus formation, was a critical driver of disease progression in influenza pneumonia with secondary bacterial infection, whereas influenza alone caused milder illness. Procoagulant platelet formation was induced via MLKL-mediated necroptosis and GSDMD-mediated pyroptosis. These platelets exacerbated pulmonary thrombosis and lung injury by inhibiting monocyte efferocytosis via complement C1qa. Platelet depletion reduced monocyte efferocytosis and worsened pneumonia, while pharmacological inhibition or platelet-specific *Gsdmd* knockout decreased procoagulant platelet levels, restored monocyte efferocytosis, and alleviated thrombotic and pulmonary injury. Mechanistically, C1qa impaired efferocytosis both by directly suppressing monocyte function and by reducing the proportion of reparative (M2-like) monocytes. Clinical relevance was confirmed by detection of MLKL/GSDMD-dependent procoagulant platelets and reduced efferocytosis receptor levels on monocytes in bronchoalveolar lavage fluid from severe pneumonia patients.

**CONCLUSIONS:** Necroptosis/pyroptosis-driven procoagulant platelets exacerbate pulmonary thrombosis by suppressing monocyte efferocytosis in a C1qa-dependent manner. These findings extend platelet-monocyte crosstalk from thrombus initiation to thrombus exacerbation, identifying modulation of the interaction between procoagulant platelets and monocyte efferocytosis as a potential therapeutic strategy for thrombus-exacerbating diseases, especially in subpopulations of patients with severe pneumonia and progressive pulmonary thrombosis.

**What Is Known?:**

1. Pulmonary thrombus exacerbation are hallmarks of severe influenza pneumonia complicated by secondary or concurrent bacterial infection, but are absent in uncomplicated mild influenza pneumonia.
2. Procoagulant platelets are key mediators of thrombotic and thromboinflammatory diseases.
3. The platelet-monocyte crosstalk via P-selectin/PSGL-1 and CD40L/CD40 initiates thrombosis, whereas the mechanisms driving progressive thrombus exacerbation remain unknown in severe pneumonia.

**What New Information Does This Article Contribute?:**

1. Necroptosis- and pyroptosis-driven elevation of procoagulant platelets mediates thrombus exacerbation, representing a key pathological feature of severe pneumonia.
2. Procoagulant platelet-impaired monocyte efferocytosis crosstalk drives thrombus exacerbation, shifting the focus from thrombus initiation to progression and offering new insight into platelet-monocyte interactions in thrombotic disease.
3. Complement C1qa may be a key mediator linking procoagulant platelets to impaired monocyte efferocytosis.

Severe influenza pneumonia with secondary bacterial infection is complicated by pulmonary thrombus exacerbation, a key contributor to respiratory failure. Although platelet-monocyte crosstalk has been implicated in thrombus initiation, whether and how this crosstalk aggravates thrombosis remains unclear. Using clinical samples, mouse models and isolated platelets challenged with influenza A virus followed by MRSA, this study shows that MLKL-mediated necroptosis and GSDMD-mediated pyroptosis promote procoagulant platelet formation. These procoagulant platelets inhibit monocyte efferocytosis via complement C1qa, thereby exacerbating pulmonary thrombosis and lung injury. Clinical relevance is confirmed by MLKL/GSDMD-dependent procoagulant platelets and reduced efferocytosis receptor levels on monocytes in BALF from severe pneumonia patients. Collectively, these findings extend platelet-monocyte crosstalk from thrombus initiation to thrombus exacerbation, identifying modulation of the interaction between procoagulant platelets and monocyte efferocytosis as a potential therapeutic strategy for thrombus-exacerbating diseases.

## INTRODUCTION

Severe influenza pneumonia has a high mortality rate and commonly occurs in the setting of secondary bacterial infection following influenza virus infection, most frequently involving *Streptococcus pneumoniae*, *Staphylococcus aureus* and *Haemophilus influenzae*. These patients often deteriorate rapidly, a core feature of which is infection-triggered coagulation cascade dysregulation, evolving into disseminated intravascular coagulation or acute respiratory distress syndrome, ultimately leading to irreversible lung injury and multiple organ failure^1^. Pathological studies have shown that extensive formation and progressive exacerbation of pulmonary thrombosis, rather than initial thrombus formation, is the central driver of disease deterioration^2^. A mouse model of 1918 H1N1 and *Streptococcus pneumoniae* coinfection displayed extensive hyaline thrombi and high tissue factor (TF) expression in the lungs, and autopsy samples from patients have verified similar changes^3^. Coagulation disorders are well documented not only in seasonal influenza, but also in severe avian influenza caused by highly pathogenic H5N1 and H7N9 infections, in which pulmonary thrombus exacerbation is frequently observed, manifesting as pulmonary thromboembolism, arterial thrombosis and hemorrhage^4^.

Platelets are anucleate megakaryocyte-derived blood cells that orchestrate inflammatory responses and thrombotic events via morphological and functional dynamics, including adhesion, activation, aggregation and intracellular signal transduction. Aberrant platelet activation and thrombocytopenia are closely linked to disease progression and poor outcomes in critical illnesses^5, 6^. Beyond canonical hemostatic functions, platelets modulate systemic coagulation homeostasis by interacting with innate and adaptive immune cells. Procoagulant platelets, a distinct platelet subpopulation characterized by surface phosphatidylserine (PS) and P-selectin (CD62P) expression, facilitate coagulation factor binding and thrombin generation, and have recently emerged as key mediators of thrombo-inflammatory disease^7^. Accumulating evidence identifies platelet-monocyte crosstalk as a core driver of hypercoagulability in severe pneumonia; for example, in severe COVID-19, activated platelets engage monocytes via P-selectin/PSGL-1 and CD40L/CD40 axes to promote aggregate formation, upregulate TF expression and enhance proinflammatory cytokine release^8, 9^. However, these interactions have been primarily studied in the context of early aggregation and activation events during thrombus initiation, whereas the mechanisms underlying thrombus exacerbation and growth, particularly in the setting of influenza-associated pneumonia with secondary bacterial infection, remain poorly understood. Current anticoagulant therapies primarily target coagulation cascade activation and thrombus formation, but their efficacy in progressive thrombus exacerbation is limited and accompanied by bleeding risks. Given that monocytes are central innate immune effectors in the lung, we hypothesize that thrombus exacerbation may be closely linked to impaired monocyte innate immune defense, particularly efferocytosis.

Efferocytosis is a multistep process where macrophages and other phagocytes recognize, engulf, and clear apoptotic cells. Defective efferocytosis is increasingly recognized as a driver of thrombotic disease progression, such as advanced atherosclerotic lesions^10^. Pulmonary Ly-6C^+^ macrophages are derived from circulating monocytes; upon pathogen challenge, these cells are recruited to the lung interstitium, differentiate into alveolar macrophages, and contribute to viral load reduction, enhanced tissue infection resistance and efferocytic function execution^11^. However, whether procoagulant platelets interact with monocyte efferocytosis to influence pulmonary thrombus exacerbation in severe pneumonia, particularly influenza virus pneumonia with secondary bacterial infection, remains unknown. This study therefore focuses on thrombus exacerbation and explores the crosstalk between procoagulant platelets and monocyte efferocytosis, as well as the underlying molecular mechanisms.

In this study, we identified elevated procoagulant platelet-driven thrombus exacerbation as a critical driver of disease progression in influenza pneumonia complicated by secondary bacterial infection, whereas influenza-only infection caused milder illness. Mechanistically, procoagulant platelet formation was induced by MLKL-mediated necroptosis and GSDMD-mediated pyroptosis, and these platelets exacerbated pulmonary thrombosis and lung injury by inhibiting monocyte efferocytosis in a complement C1qa-dependent manner. Our findings expand the understanding of platelet-monocyte crosstalk from thrombus initiation to thrombus exacerbation in pneumonia-associated coagulopathy and identify procoagulant platelets and monocyte efferocytosis as potential therapeutic targets for severe viral-bacterial pneumonia and related thrombotic disorders.

## METHODS

### Mice and ethics

WT C57BL/6J mice (6-8 weeks old) were purchased fromVituvilua Biotechnology Co., Ltd. (Guang Zhou, China). Animals were randomly assigned to control or experimental groups using a computer-generated randomisation sequence (GraphPad QuickCalcs). All mice were housed in specific pathogen-free facilities with free access to food and water. All experimental protocols were approved by the Animal Care and Use Committee of Guangzhou National Laboratory (Permit number: GZLAB-AUCP-2024-11-A07). *Nlrp3*^-/-^, *Gsdmd*^-/-^ and platelet-specific *Gsdmd* knockout (*Gsdmd^flox/flox^* PF4-Cre) mice were gifted by Prof. Jing-lin Wang and Yuan Yuan from the Academy of Military Medical Sciences. Platelet-specific *Gsdmd* knockout mice (*Gsdmd^flox/flox^* PF4-Cre) were generated by crossing *Gsdmd flox* and PF4-Cre mice. Mice were genotyped using primers P1 (5′-CCAAGTCCTACTGTTTCTCACTC -3′) and P2 (5′-TGCACAGTCAGCAGGTT-3′) for PF4-Cre; P3 (5 ′ -TAGAAACAACAGCAGGTTGGGAA-3 ′) and P4 (5 ′ -TGGGGTATGACAAGTTCAAACACG-3 ′) for *Gsdmd flox*. All procedures were conducted under BSL-2 conditions.

### Clinical Sample of Severe Pneumonia Collection and Analysis

The healthy donors and severe pneumonia patients (caused by multiple bacterial or viral pathogens) who supplied blood samples for this study were provided with written informed consent in accordance with the Declaration of Helsinki (Health Control, n=15; Severe Pneumonia, n=28). Approval was obtained from the Institutional Medical Ethics Committee of the Affiliated Panyu Central Hospital of Guangzhou Medical University (Permit number: PYRC-2025-209-01). Severe pneumonia patients were defined according to the 2007 Infectious Diseases Society of America/American Thoracic Society (IDSA/ATS) consensus criteria, requiring the presence of at least one major criterion or three or more minor criteria [citation to IDSA/ATS guideline]. Major Criteria: (1) Need for endotracheal intubation and mechanical ventilation; (2) Septic shock requiring vasopressors after adequate fluid resuscitation. Minor Criteria: (1) Respiratory rate ≥ 30 breaths/min; (2) PaO_2_/FiO_2_ ratio ≤250 mmHg; (3) Multilobar infiltrates on chest imaging; (4) Confusion or disorientation; (5) Blood urea nitrogen (BUN) ≥7.14 mmol/L (≥20 mg/dL); (6) Systolic blood pressure <90 mmHg; (7) Need for aggressive fluid resuscitation.

### Pathogens and infections

Influenza A virus (A/PR/8/34; Sinai strain) was used to construct a severe pneumonia model and in *vitro* analysis of influenza virus induced platelet thrombus formation. This virus was propagated in 10-day-old embryonated chicken eggs. The methicillin-resistant *Staphylococcus aureus* (MRSA) strain USA300 (BAA-1556) were purchased from the American Type Culture Collection. The MRSA strain was cultured and propagated through Luria-Bertani plate or medium.

### Cell lines

The cell line THP-1 and RAW264.7 cell was sourced from the ATCC. Cells were maintained at 37°C, 5% CO_2_ in RPMI 1640 supplemented with 10% heat-inactivated FBS and 1% penicillin-streptomycin. All procedures were conducted under BSL-2 conditions.

### Establishment of a mouse model of IAV and MRSA infected pneumonia

Constructing a severe pneumonia model in mice by intra-nasal infection with IAV and MRSA. 7-week-old mice were intranasally infected with 1 LD_50_ IAV. Three days later, mice were intranasally administered vehicle (PBS) or 10^8^ CFU of MRSA and were euthanized 6 hours later. The blood, bronchoalveolar lavage fluid (BALF) and lung tissue of mice were collected. *Gsdmd^flox/flox^* PF4-Cre and *Gsdmd^flox/flox^* mice were also sequentially infected with IAV and MRSA as described above for analyzing the role of platelet GSDMD in severe pneumonia caused by the sequential infections. The lung index (the ratio of lung tissue weight to body weight), the number of cells in BALF and daily weight changes were recorded. Severe pneumonia in mice is defined as meeting at least two of the following three criteria: (1) Body weight loss >5%, obvious tachypnea (markedly increased respiratory rate) or dyspnea, markedly reduced activity, huddling, piloerection or hunched posture; (2) Pulmonary inflammatory infiltration covering >50% of the lung area, hyaline thrombus formation in >10% of pulmonary vessels and a histopathological score >5; (3) Neutrophil percentage in bronchoalveolar lavage fluid (BALF) >50%, and levels of IL-1β and TNF-α ≥2-fold higher than those in the normal control group. All procedures were performed under light isoflurane anesthesia to minimise pain and distress during virus inoculation, bacterial challenge, and sample collection. Mice were monitored daily for body weight, clinical signs, and behaviour. Humane endpoints were predefined: animals with >25% body weight loss, severe lethargy, or moribund condition were euthanised immediately by CO_2_ inhalation followed by cervical dislocation.

### Platelet depletion mouse model

Platelets were depleted from C57BL/6J mice as previously described^12^. Briefly, mice were injected intra-peritoneally with anti-platelet glycoprotein Ib beta chain (GPIb, CD42b, cat#R300, Emfret Analytics) or anti-immunoglobulin-G (IgG, cat#C301, Emfret Analytics) antibodies, at 4 μg of antibody/gram body weight of mouse. At 48 h post ablation, mice were infected with influenza A virus. Subsequently, the mice continued to undergo platelet elimination two days after infection with IAV. 24 hours later, MRSA was administered via nasal drip to establish a platelet depletion model in mice with severe pneumonia induced by sequential infections of IAV and MRSA.

### Stimulation of mouse platelets *in vitro* with IAV and MRSA

The mouse blood was collected using sodium citrate anticoagulant, followed by centrifugation at 200g for 15 minutes to obtain plasma rich platelets (PRP). All stimulation experiments were conducted on PRP to mimic the in *vivo* environment in humans. That is, IAV (at a ratio of 1 PFU:100 platelets) was first used to stimulate PRP for 1 hour, followed by MRSA addition to continue stimulating for another 15 minutes. Subsequently, the PRP was centrifuged at 500g for 10 minutes after stimulation, the highly platelet like precipitates were used for immunofluorescence (IF) and Western-blot (WB) analysis. Whatever, before stimulation, 10μM Necrosulfonamide (NSA, GSDMD and MLKL inhibitor, HY-100573, MedChemExpress), as well as 50μM GSK-872 (PIPK3 inhibitor, HY-101872, MedChemExpress) were added 30 minutes in advance. And, PRP from *Nlrp3*^-/-^, *Gsdmd*^-/-^ and WT mice, as well as PRP from *Gsdmd^flox/flox^* PF4-Cre and *Gsdmd^flox/flox^* mice, was also isolated and subjected to stimulation with IAV and MRSA. The changes in platelet coagulation factors and inflammatory cytokines were analyzed.

### Immunoblot of platelets

Platelet proteins were extracted in the presence of lysis buffer (including PMSF and phosphatase inhibitor, at a final concentration of 1mM). Protein quantitation was performed using the bicinchoninic acid (BCA) method (P0399M, Beyotime). Proteins were separated on 10% polyacrylamide SDS-PAGE gels (PG112, Epizyme), with a load of 20-30μg, and then transferred onto a polyvinylidene difluoride (PVDF) membrane (IPVH00010, Millipore) via electro-transfer. The membrane was blocked with 5% nonfat dry milk in TBS with 0.1% Tween 20 (TBST) for 2.5 hours. Primary antibodies were diluted in immunoblotting (abs954, Absin) and incubated overnight at 4°C. The membrane was then incubated with HRP-labeled Goat anti-rabbit IgG or HRP-labeled Goat anti-mouse IgG (1:5000 in 5% nonfat dry milk in TBST) for 1 hour at room temperature. Blots were detected using the ECL luminescent reagent (P90719, Millipore) and imaged using an ImageQuant LAS4000 system (Boston, GE Healthcare). The specific antibodies are as follows, anti-p-RIPK1 (1:1000, #AF7377, Affinity), anti-RIPK1 (1:1000, #DF2642, Affinity), anti-p-RIPK3 (1:1000, #AF7443, Affinity), anti-p-RIPK3 (1:1000, #DF10141, Affinity), anti-p-MLKL (1:1000, #37333, CST), anti-MLKL (1:1000, #37705, CST), anti-NLRP3 (1:1500, MAB7578, R&D Systems) and anti-GSDMD (1:200, ab219800, abcam).

### Stimulation of macrophages by platelet supernatant and detection of efferocytic function

After mouse PRP being stimulated by IAV and MRSA, the supernatants of the PRP were collected and the monocyte RAW264.7 cells were stimulated at a ratio of 25μL/10^5^ cells. In parallel, THP-1 cells were subjected to stimulation with PRP supernatant obtained from patients with severe pneumonia caused by multiple bacterial or viral pathogens. After 24 hours of treatment, these cells were collected and their efferocytic function was observed following previous reports (ab234053, abcam)^13^. The percentage of reparative monocytes (CD206^+^ in RAW264.7 cells and CX3CR1 in THP-1 cells) was also performed using flow cytometry. In addition, to analyze the effect of C1qa on monocyte efferocytosis, 30μg/mL C1qa (CSB-EP003637MOa0, CUSABIO) was also added along with platelet supernatant.

### Flow cytometry detection of immune cells and platelets

Flow cytometry was used to detect immune cells (neutrophils, monocytes and macrophages, etc.) as well as platelet associated CD62P, IL-1β, TNF-α, CD40L and TF in BALF of mice with severe pneumonia. At the same time, in *vitro* platelets were stimulated by IAV and MRSA, and platelet associated activation, inflammation and coagulation molecules were detected by flow cytometry.

Platelet related activation and coagulation in clinical severe pneumonia samples were also detected by flow cytometry. And the phenotype analysis related to monocytes were detected using flow cytometry. Please refer to the attachment for specific operation methods.

### Immunofluorescence detection of platelet and monocyte related changes in lung tissue and blood in mice and patients

The lung tissues of mice with IAV and MRSA infected pneumonia and control were fixed with 4% paraformaldehyde and sectioned in paraffin. After dewaxing and antigen repair, lung tissue was stained with corresponding antibodies for anti-CD41 (1:50, sc-365938, santa cruz biotechnology), anti-Ly-6C (1:200, ab317272, abcam), anti-Fibrin (1:100, clone 59D8, sigma), anti-FXIIIa (1:100, ab1834, abcam), anti-TF (1:50, 28005, Proteintech), anti-Mertk (1:50, sc-365499, santa cruz biotechnology), anti-GSDMD (1:200, abcam, ab219800), anti-p-MLKL (1:200, #37333, CST) and anti-C1qa (1:200, CSB-PA003637GA01HU, CUSABIO).

BALF from patients with severe pneumonia was collected, concentrated by centrifugation, fixed, and then stained with corresponding antibodies for anti-CD41 (1:50, sc-365938, santa cruz biotechnology), anti-Ly-6C (1:200, ab317272, abcam), anti-TF (1:50, 28005, Proteintech), anti-Mertk (1:100, sc-365499, santa cruz biotechnology) and anti-CX3CR1 (1:100, 29819-1-AP, Proteintech).

Blood samples from patients with severe pneumonia caused by multiple bacterial or viral pathogens were collected and the platelets were separated. And the platelets were stimulated with IAV and MRSA in *vitro* using antibodies anti-CD41(1:50, sc-365938, santa cruz biotechnology), anti-p-RIPK1 (1:200, #AF7377, Affinity), anti-p-MLKL (1:1000, #37333, CST), anti-GSDMD (1:200, ab219800, abcam), anti-CD40L (1:200, DF2301, Affinity), anti-TF (1:50, 28005, Proteintech) and anti-C1qa labeling (1:200, CSB-PA003637GA01HU,CUSABIO) to complete platelet related death, coagulation factor and complement detection.

After overnight incubation, the secondary antibodies used were Alexa Fluor 488-conjugated anti-mouse IgG antibody (1:1000, ab150113, Abcam) and Alexa Fluor 647-conjugated anti-rabbit IgG antibody (1:1000, ab150075, Abcam). Imaging was performed using a ZEISS LSM 900 confocal microscopy with a 60×immersion lens.

### ELISA

The levels of TAT (CSB-E08433m,CUSABIO), TF (JL10803, Shanghai Jianglai Biotechnology Co., Ltd), sCD62P (CSB-E04709m,CUSABIO), PF4 (MM-0076M1, MEIMIAN), HMGB1 (MM-44107M1, MEIMIAN), TM (JL11551, Shanghai Jianglai Biotechnology Co., Ltd), t-PAI-c (MM-46769M1, MEIMIAN), IL-1β (CSB-E08054m, CUSABIO), IL-6 (CSB-E04639m, CUSABIO), TNF-α (CSB-E04741m, CUSABIO) and C1qa (CSB-EL003637MO, CUSABIO) in plasma and BALF of mouse were measured using ELISA kits, as per the manufacturer’s instructions.

### Observation of platelet morphology under transmission electron microscope (TEM)

In order to clarify the phenotype of platelet morphology, platelet in patients with severe pneumonia, as well as platelets underwent IAV and MRSA stimulation were fixation. The ultrastructural sections were examined with the TEM (Japan Electron Optics Laboratory, JEM-1400), followed captured with an Advantage CCD camera (MORADA, EMSIS) using RADIUS AII 2.2 (build 21230) software.

### Statistical analysis

The data were plotted and analyzed using GraphPad Prism 10.1.2. software. Statistical significance was assessed using the unpaired two-tailed Student’s *t*-test, one-way or two-way analysis of variance followed by a *post hoc* Newman-Keuls test, where appropriate, and was considered significant at *P*<0.05. And correlation analysis was statistically analyzed using Pearson’s correlation analysis. Normality of data distribution was assessed using the Shapiro-Wilk test, and homogeneity of variance was assessed using the Brown-Forsythe test. For data that met these assumptions, parametric tests (unpaired two-tailed Student’s t-test or ANOVA) were used. For data that did not meet the normality assumption, non-parametric equivalents (Mann-Whitney U test or Kruskal-Wallis test) were used. If variance homogeneity was not met, Welch’s correction was applied.

## RESULTS

### Elevated procoagulant platelets mediate pulmonary thrombus exacerbation correlated with higher severity induced by secondary bacterial infection

In cases of human severe pneumonia caused by influenza virus infection, bacterial co-infection or secondary infection commonly occurs, and these patients frequently present with pulmonary microangiopathy and pulmonary thrombosis, accompanied by more severe disease progression and higher mortality^1, 2^. To characterize the pathological role of platelets and pulmonary thrombosis played in severe influenza pneumonia, we constructed a mouse model of sequential infections with influenza A virus (PR8: A/Puerto Rico/8/1934, IAV) and methicillin-resistant *Staphylococcus aureus* (MRSA) **(Figure 1A)**. Unlike influenza virus infection alone, influenza virus with secondary bacterial infection caused further reductions in lung index, increases in cell counts in BALF and greater body weight loss in mice **(Figure 1B)**. To evaluate the correlation between platelets and pulmonary thrombosis in severe pneumonia, lung tissues were examined by hematoxylin and eosin (H&E) staining and immunofluorescence (IF) staining. The results showed that influenza virus with secondary bacterial infection led to significantly increased 15% hyaline thrombi **(black arrow was shown in Figure 1C)**, Fibrin deposition **(yellow arrow in Figure 1C)**, 50% inflammatory infiltration and high tissue factor (TF)-expressing platelets **(white arrow in Figure 1C) (Figures 1C, 1D, S1A and S1B)**. Flow cytometric analysis of platelets in BALF further demonstrated that influenza virus with secondary bacterial infection significantly recruited more platelets into the mouse lungs, and these platelets exhibited significantly upregulated expression of CD62P, CD40L, IL-1β and TNF-α **(Figure 1E)**. A significant increase in procoagulant platelets, characterized as CD62P^+^ and TF^+^, was confirmed in mice with secondary bacterial infection following influenza virus challenge. There were significantly elevated levels of coagulation markers (TF and thrombin-antithrombin complex [TAT]), platelet-activating factors (PF4 and sCD62P), the damage-associated molecule HMGB1 and pro-inflammatory cytokines (IL-1β and TNF-α) in BALF of influenza virus with secondary bacterial infection **(Figures 1F, S1C-S1E)**. Further correlation analysis indicated that hyaline thrombi and platelets with high expression of TF, CD40L and IL-1β were significantly positively correlated with pathological damage **(Figure 1G and S1F)**. These findings suggest that severe lung injury induced by influenza virus with secondary bacterial infection is closely associated with elevated procoagulant platelet-mediated pulmonary thrombus exacerbation.

**Figure 1.**
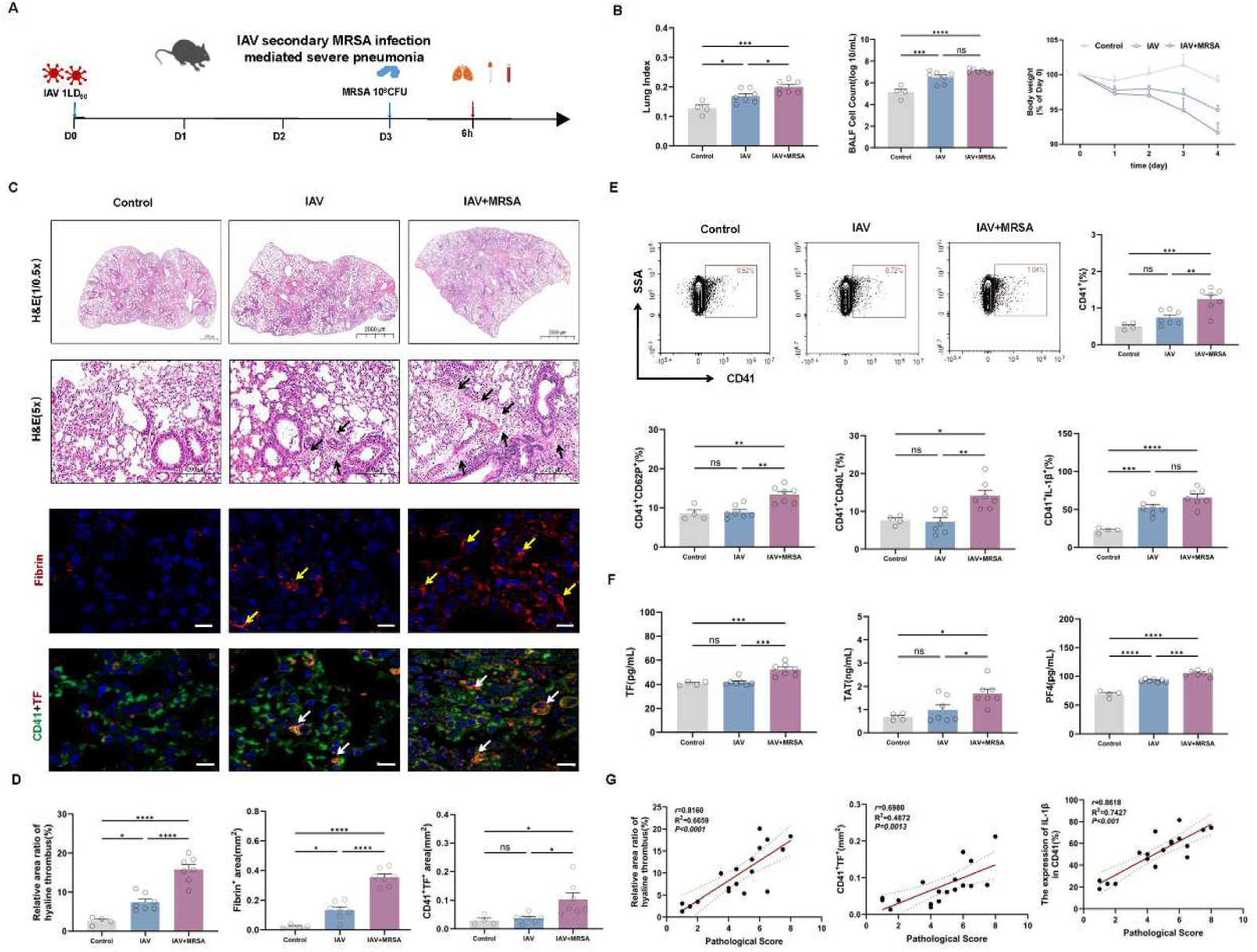
Elevated procoagulant platelets mediate pulmonary thrombus exacerbation correlated with higher severity induced by secondary bacterial infection. **(A)** The experimental scheme:7-week-old mice were intranasal infected with 1 LD_50_ HIN1 influenza virus (PR8: A/Puerto Rico/8/1934). Three days later, mice were intranasally administered vehicle (PBS) or 10^8^ CFU of MRSA and were euthanized 6 hours later (Control, n=4; IAV, n=7; IAV+MRSA, n=7). **(B)** The lung index (the ratio of lung tissue weight to body weight), the number of cells in bronchoalveolar lavage fluid (BALF), and daily weight changes were recorded. **(C and D)** The pathological damage of mouse lung tissue was completed through H&E staining and analysis, bar, 2000, 1000 or 200 μm. The black arrows indicated hyaline thrombi. The expression of Fibrin and CD41^+^TF^+^ in lung tissue was measured by IF, respectively, and semi quantitative analysis was performed using Image J, bar, 10 μm. The yellow and white arrows indicated Fibrin and CD41^+^TF^+^. **(E)** Flow cytometry detection of platelet counts in BALF and expression of CD62P, IL-1β, TNF-α and CD40L derived from platelets. **(F)** Tissue factor (TF), Thrombin and antithrombin complexes (TAT) and platelet factor 4 (PF4) were measured. **(G)** The relatively percentage of lung hyaline thrombic, CD41^+^TF^+^ in lungs, as well as platelet derived IL-1β in BALF were strongly positively correlated with the degree of injury. Data were presented as mean ± SEM and statistical analysis was done using one-way ANOVA **(B and D-F)** or Pearson’s correlation analysis **(G)**, ns *P*>0.05, * *P* <0.05, ** *P* <0.01, *** *P* <0.001, **** *P* <0.0001.

### Platelet pyroptosis and necroptosis drives procoagulant platelets in influenza virus with secondary bacterial infection

To further elucidate the role of procoagulant platelet generation in severe pneumonia, we also established an in *vitro* model of platelets with secondary MRSA stimulation following IAV stimulation **(Figure 2A)**. The results showed that influenza virus alone increased the PS^+^ population. In contrast, influenza virus with secondary bacterial stimulation significantly elevated PI^+^ (necrotic) platelets **(Figure 2B)**. Consistent with the results of the in *vivo* study, significantly elevated platelet activation, inflammation and procoagulant platelets, defined as PS^+^, CD62P^+^ and TF^+^ were observed following secondary bacterial stimulation after influenza virus infection. **(Figures 2B, 2C and 2E)**. Previous studies have shown that programmed platelet death is closely linked to platelet coagulatory activity and inflammatory responses^14, 15^. Therefore, we further characterized the programmed cell death modes of platelets, including necroptosis and pyroptosis **(Figure 2B)**. The results showed that secondary bacterial stimulation following influenza virus stimulation significantly upregulated the NLRP3/activated caspase-1/GSDMD-N and p-RIPK1/p-RIPK3/p-MLKL protein expression involved both pyroptotic and necroptotic pathways **(Figures 2D-F)**. Immunofluorescence and transmission electron microscopy (TEM) of platelet further confirmed morphological changes: influenza virus alone induced platelet shrinkage **(as shown by red arrows in Figure 2H)**, whereas influenza virus secondary bacterial stimulation caused platelet enlargement **(yellow arrows in Figure 2H and white arrows in Figure 2G)**, membrane swelling **(white arrows in Figure 2G)**, rupture and cytoplasmic leakage **(blue arrows in Figure 2G)**, consistent with the morphology of necroptosis and pyroptosis **(Figures 2G and 2H)**. Furthermore, platelets in the lungs of mice with secondary bacterial infection also exhibited elevated levels of GSDMD and p-MLKL **(Figure 2I)**. These findings indicate that influenza virus infection with secondary bacterial infection induces pyroptotic, necroptotic and procoagulant platelets.

**Figure 2.**
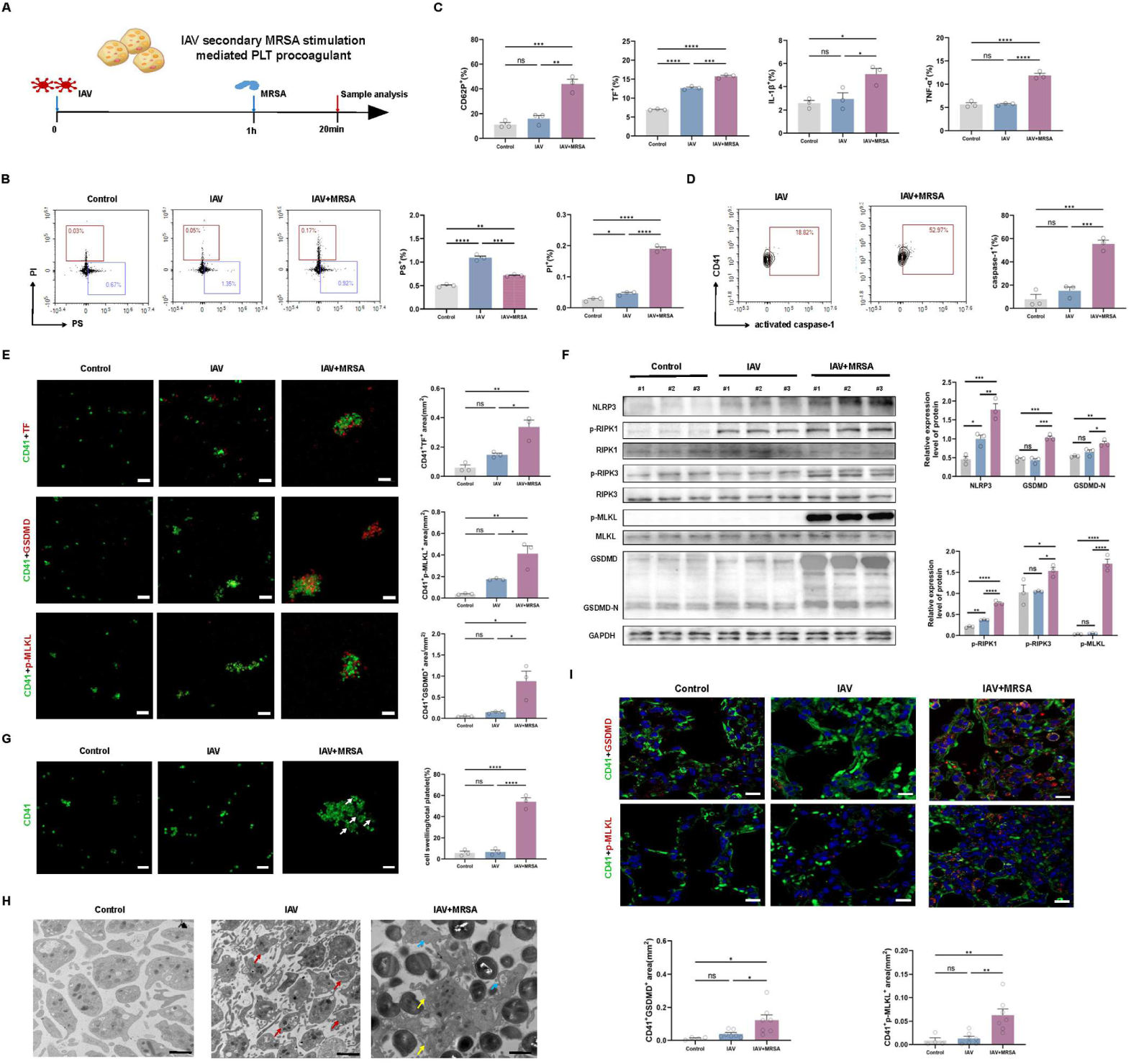
Platelet pyroptosis and necroptosis drives procoagulant platelets. (A) The experimental scheme: MRSA infection secondary to IAV activates platelets and induces platelet procoagulant activity and inflammation. **(B)** Platelet death was detected through staining PS and PI (n=3). **(C and D)** TF, CD62P, caspase-1, IL-1β and TNF-α derived from platelets were determined using flow cytometry (n=3). **(E)** Platelet related TF, GSDMD and p-MLKL were detected by IF (n=3). Scale bars, 10 μm. **(F)** Influenza virus secondary bacterial stimulation of platelets leaded to increased expression of platelet necroptosis related proteins (p-RIPK1-p-RIPK3-p-MLKL) and pyroptosis related proteins (NLRP3-GSDMD) (n=3). **(G)** Immunofluorescence staining revealed the morphology of platelet. The white arrow represented broken platelets. Scale bars, 10 μm (n=3). **(H)** Transmission electron microscopy showed that platelet morphology when platelet stimulated by IAV or IAV secondary MRSA infection. IAV induced platelet shrinkage, but IAV secondary MRSA stimulation led to platelet enlargement, cell membrane damage, structural damage and vesicle secretion. The red, yellow and blue arrows respectively represented shrunken, swollen platelets and platelet-derived microvesicles. Scale bars, 1 μm. **(I) Necrotic and pyroptosis platelets in mice with severe pneumonia.** The relative expression of GSDMD and p-MLKL derived from platelets in lung tissue (Control, n=4; IAV, n=7; IAV+MRSA, n=7). Data were presented as mean ± SEM and statistical analysis was done using one-way ANOVA (**B-G and I)**, ns *P>*0.05, \**P*<0.05, \*\**P*<0.01, \*\*\**P*<0.001, \*\*\*\**P*<0.0001.

To establish a causal relationship between pyroptosis or necrosis signals and procoagulant activity, GSK 872 (a RIPK3 inhibitor) and NSA (GSDMD and MLKL inhibitors) were used to pharmacologically inhibit platelets stimulated by secondary bacterial stimulation. Early administration of GSK-872 and NSA suppressed platelet lytic death, accompanied by increased platelet counts and decreased levels of fibrinogen, TF, CD62P, IL-1β and TNF-α **(Figures S2A-S2C)**. We further isolated platelets from WT, *Gsdmd*^-/-^ and *Nlrp3*^-/-^ mice and subjected them to sequential IAV and MRSA treatment. Compared with WT platelets, platelets derived from *Gsdmd*^-/-^ and *Nlrp3*^-/-^ mice exhibited markedly reduced expression of CD62P, CD40L, TF, IL-1β and TNF-α, with the most pronounced attenuation in *Nlrp3*^-/-^ platelets **(Figures S2D and S2E)**. To further corroborate these findings, we isolated platelets from *Gsdmd^flox/flox^* PF4-Cre and control mice and sequentially challenged them with IAV and MRSA *ex vivo* **(Figure S2F)**. Control platelets upon pathogen stimulation exhibited robust upregulation of GSDMD, CD40L, CD62P, TNF-α, IL-1β and TF, along with elevated supernatant fibrinogen. In contrast, platelets from *Gsdmd^flox/flox^*PF4-Cre mice with pathogen stimulation showed significantly declined responses, with IL-1β displaying the most pronounced reduction **(Figures S2G-S2I)**. These results confirm that pyroptosis and necroptosis synergistically drives procoagulant platelet formation.

### Procoagulant platelets inhibit monocyte efferocytosis during influenza infection complicated with secondary bacterial infection

Platelets not only directly participate in coagulation and inflammatory responses but also exert these effects through their interactions with monocytes and neutrophils. Accumulating evidence identifies platelet-monocyte crosstalk as a core driver of hypercoagulability in severe pneumonia^8^. We therefore investigated platelet-monocyte interactions to thrombus exacerbation in a murine model of influenza virus infection complicated by secondary bacterial challenge. By staining and classifying distinct immune cell populations in BALF from mice with severe pneumonia induced by the dual infection, we observed that influenza virus alone significantly reduced the number of lung resident macrophages (CD11b^-^F4/80^+^Siglec-F^+^) while increasing infiltrating monocytes (F4/80^-^CD11b^+^Ly-6C^+^). Notably, this monocyte population was markedly depleted following secondary bacterial challenge **(Figures 3A and 3B)**. These infiltrating monocytes expressed high levels of CX3CR1 and Mertk **(Figures 3B and 3C)**, both of which are critical receptors for efferocytosis, the phagocytic clearance of apoptotic cells. Specifically, CX3CR1 mediates the "find me" signal that guides monocytes to apoptotic targets, whereas Mertk coordinates the subsequent "eat me" phase for engulfment. Furthermore, the number of monocytes with high CX3CR1 expression correlated negatively with disease severity **(Figure 3B)**. Collectively, these results demonstrate that severe influenza pneumonia with secondary bacterial infection is associated with reduced levels of efferocytic receptors on monocytes.

**Figure 3.**
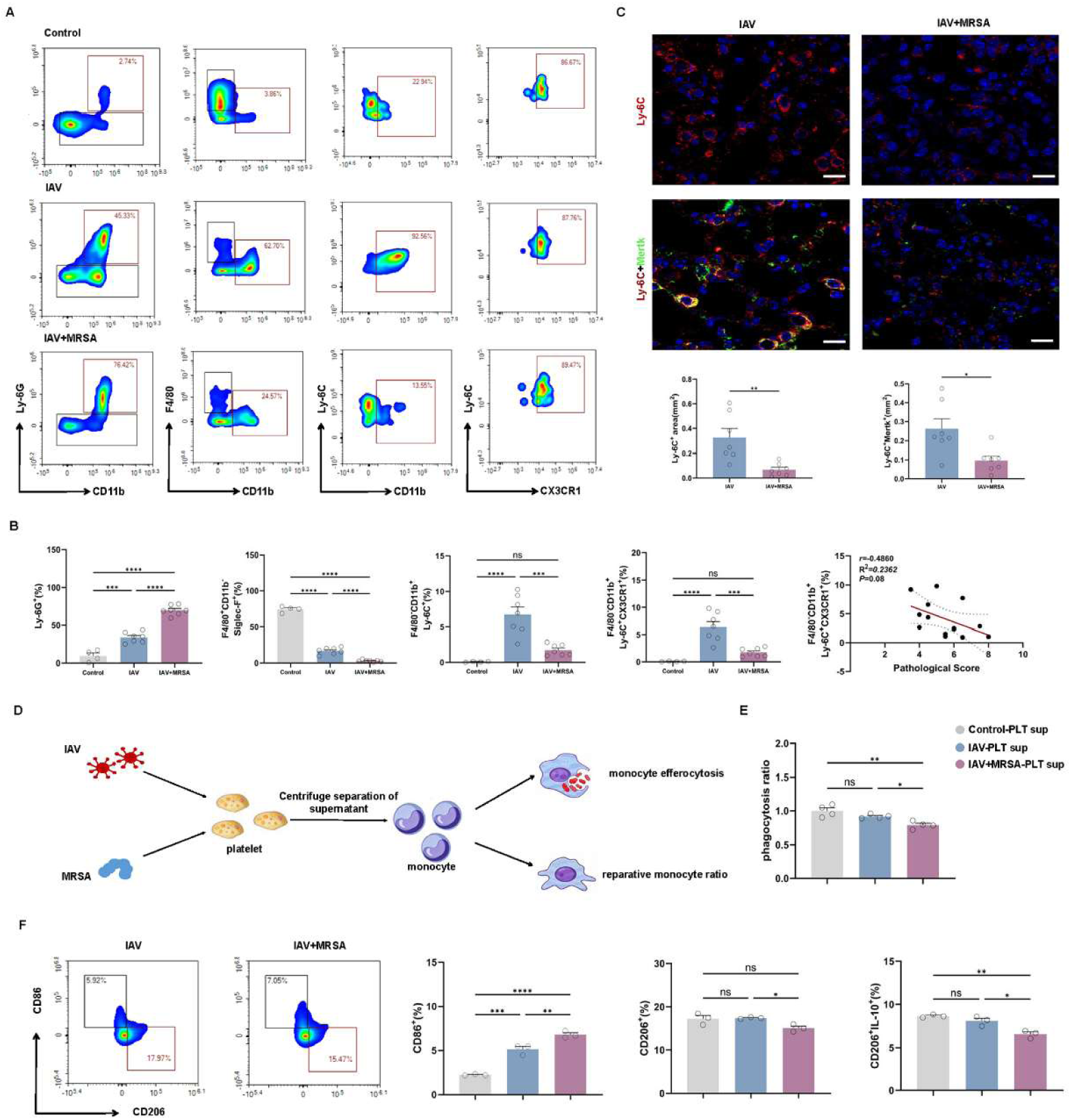
Procoagulant platelets suppress monocyte efferocytosis in influenza virus with secondary bacterial infection. **(A-D) Impaired efferocytosis receptor expression on monocytes in severe pneumonia caused by influenza virus and secondary bacterial infection. (A and B)** Flow cytometry was used to detect the proportion of neutrophils, macrophages and monocytes in BALF based on distinct surface markers. Ly-6G^+^, CD11b^-^F4/80^+^Siglec-F^+^, CD11b^+^F4/80^-^Ly-6C^+^ and CD11b^+^F4/80^-^Ly-6C^+^CX3CR1^+^ represent neutrophils, storage macrophages, monocyte and efferocytic monocytes respectively (Control, n=4; IAV, n=7; IAV+MRSA, n=7). Influenza virus infection significantly increased monocyte, while secondary bacterial infection significantly reduced these cells. Meanwhile, these cells highly express CX3CR1. Pearson’s correlation analysis indicates that the quantity of CD11b^+^F4/80^-^Ly-6C^+^CX3CR1^+^ was negatively correlated with the pathological score of mice. **(C)** Immunofluorescence was used to detect the localization and expression of monocytes (Ly-6C, red) and efferocytosis receptor (Mertk, green) in lung tissue. **(D-F) Supernatant from platelets exposed to influenza virus and secondary bacterial infection inhibits monocyte efferocytosis and reduces the proportion of reparative monocytes**. **(D)** Schematic: RAW264.7 cells were co-cultured with supernatant from platelets that had been secondarily challenged with IAV and subsequently MRSA, and the efferocytosis and reparative monocyte proportion were evaluated. **(E)** After different stimuli of platelet supernatant were applied to RAW264.7 cells, efferocytosis was performed (n=4). **(F)** The proportion of reparative monocytes (CD206^+^) and the IL-10 secreted by these monocytes were calculated (n=3). Data were presented as mean ± SEM and statistical analysis was done using one-way ANOVA **(B, E and F)** or Unpaired *t* test **(C)**, ns *P>*0.05, \**P*<0.05, \*\**P*<0.01, \*\*\**P*<0.001, \*\*\*\**P*<0.0001.

To directly characterize the inhibitory effect of procoagulant platelets on monocyte efferocytosis, we exposed monocytes in *vitro* to supernatants from platelets subjected to various stimuli, and then assessed efferocytic function and the proportion of reparative monocytes **(Figure 3D)**. Supernatant from shrunken platelets induced by influenza virus alone did not reduce monocyte efferocytosis and decrease the frequency of reparative (CD206^+^) monocytes. In contrast, supernatant from platelets that underwent necrosis and pyroptosis following dual pathogen stimulation significantly impaired efferocytosis, lowered the reparative monocyte proportion and diminished IL-10 expression in these cells **(Figures 3E and 3F)**. Notably, when platelets were pretreated with GSK-872 and NSA inhibitors before sequential influenza and MRSA challenge, the resulting supernatant rescued monocyte efferocytosis and restored the reparative monocyte frequency **(Figures S3A and S3B)**. Similarly, supernatant from platelets isolated from *Gsdmd^flox/flox^* PF4-Cre mice, subjected to the same sequential stimulations, failed to impair efferocytosis and reduce the reparative monocyte population **(Figures S3C and S3D)**. Collectively, these findings indicate that supernatant from procoagulant platelets driven by necroptosis and pyroptosis is associated with impaired monocyte efferocytosis and a reduced pool of reparative monocytes, which may represent a contributing mechanism to pulmonary thrombus exacerbation in severe pneumonia with secondary bacterial infection.

### Thrombocytopenia impairs monocyte efferocytosis and aggravates pulmonary thrombosis in influenza virus–infected mice

To clarify the association between platelets and monocyte efferocytosis on the exacerbation of pulmonary thrombosis in *vivo*, we established thrombocytopenic mouse models in wild-type C57BL/6J mice using an anti-glycoprotein Ibα (GPIbα) antibody, as previously described^12^, and evaluated monocyte efferocytosis, pulmonary thrombosis and lung injury following influenza virus infection and secondary bacterial challenge. After anti-GPIbα treatment, platelet counts were decreased in mice **(Figure 4A)**. Compared with anti-IgG controls, anti-GPIbα-treated infected mice exhibited more severe lung injury, characterized by enhanced hyaline thrombus formation and inflammatory cell infiltration **(black and red arrows in Figure 4B)**, along with significantly increased lung index and viral loads **(Figures 4B and S4A)**. These mice also displayed higher pulmonary levels of Fibrin, activated factor XIII (FXIIIa), and TF **(Figure 4C)**, as well as elevated concentrations of TAT, thrombomodulin (TM), tissue plasminogen activator-plasminogen activator inhibitor-1 complex (t-PAI-C) in BALF and plasma, and increased IL-6 and IL-1β in BALF **(Figure 4D, S4B and S4C)**. These data demonstrated that thrombocytopenia exacerbated pulmonary thrombosis and inflammation. We further examined monocyte efferocytosis and its receptor expression in this context. Compared with anti-IgG mice, thrombocytopenic mice showed a pronounced reduction in F4/80^-^CD11b^+^Ly-6C^+^, CX3CR1^+^Ly-6C^+^, and Mertk^+^Ly-6C^+^ monocyte populations, particularly after influenza virus infection **(Figures 4E-4G and S4D)**. Collectively, these findings suggest that thrombocytopenia is associated with exacerbated pulmonary thrombosis and lung injury in pneumonia, likely through suppression of monocyte efferocytosis.

**Figure 4.**
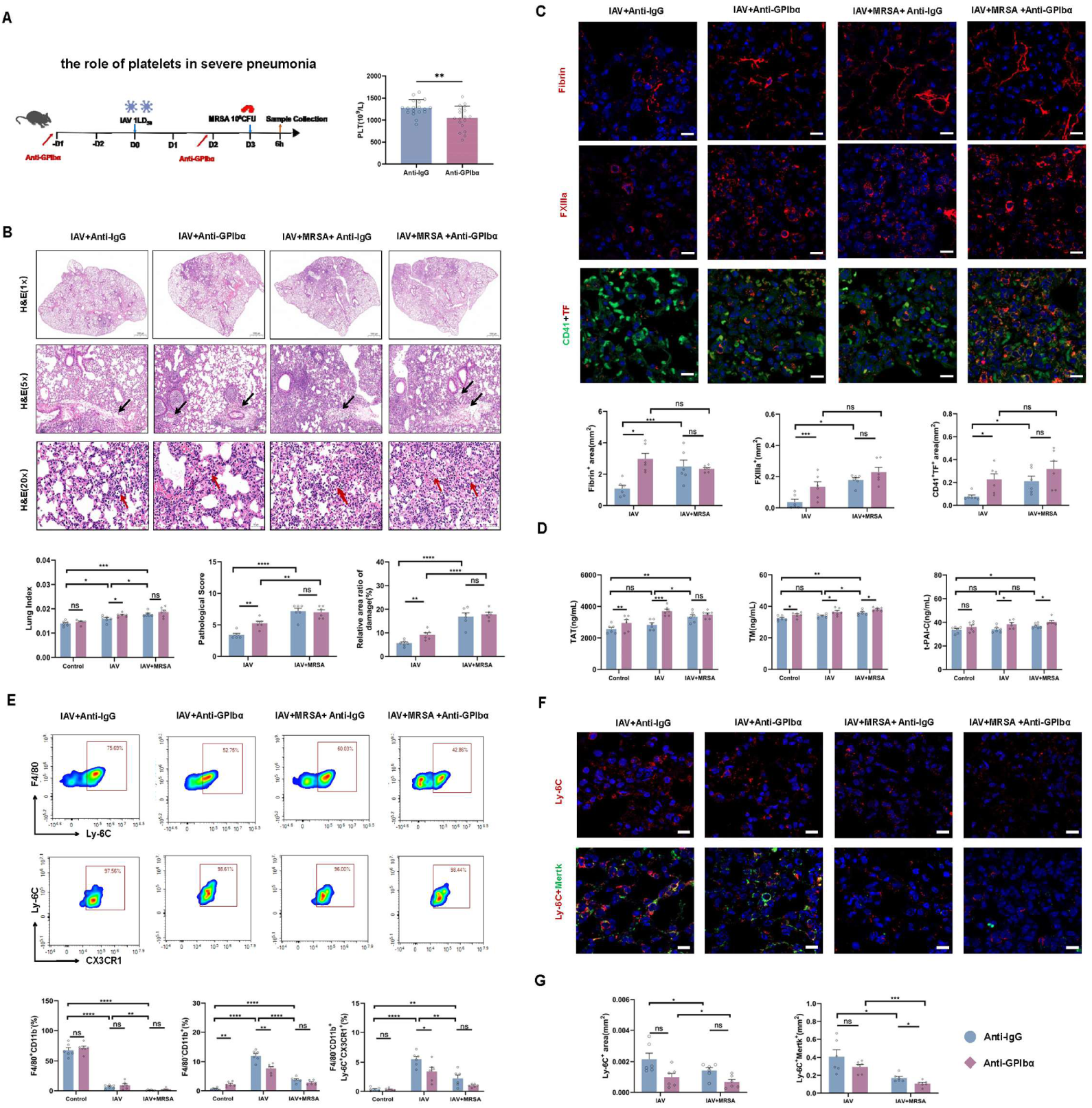
Thrombocytopenia impaires monocyte efferocytosis and aggravates pulmonary thrombosis in influenza virus–infected mice. (A) Schematic: Using GPIbα antibody to block platelets for observing the role of platelets in severe pneumonia and to evaluate monocyte efferocytosis and thrombus exacerbation. Simply put, anti-GPIbα and anti-IgG was administered two days before IAV infection, followed by IAV infection in mice, and these groups respectively were referred to as IAV + Anti-GPIbα and IAV + Anti-IgG; Two days before and after IAV infection, anti-GPIbα and anti-IgG was administered, and 12 hours later, mice were intranasal infected with MRSA. The two group respectively were referred to as IAV+MRSA + Anti-GPIbα and IAV+MRSA + Anti-IgG. The number of platelets in mice was also determined after anti-GPIbα or anti-IgG treatment (n=6). **(B)** The pathological damage of mice (transparent thrombus and inflammatory infiltration) was presented by H&E staining and semi quantitative calculation using Image J (bar, 1000, 200 or 50 μm). The black and red arrows indicate hyaline thrombi and inflammatory cell infiltration, respectively. The lung index of mice was calculated by comparing the weight of lung tissue to the body weight of mice. **(C)** The expression of thrombotic factors (Fibrin, FXIIIa and TF) in mice after Anti-GPIbα antibody intervention was measured by Immunofluorescence analysis. Lung tissues were stained with nucleus for DAPI (blue), anti-Fibrin, FXIIIa and TF (red) and CD41 for platelets (green). Scale bars, 10 μm. **(D)** ELISA determination of thrombin antithrombin complex (TAT), thrombomodulin (TM) and tissue plasminogen activator-plasminogen activator inhibitor-1 complex (t-PAI-C) in mouse BALF. **(E)** BALF monocytes with efferocytosis receptor were analyzed via CD11b^+^F4/80^-^Ly-6C^+^CX3CR1^+^ staining. **(F)** Lung cells co-expressing Ly-6C and Mertk were examined by IF staining. Data were presented as mean ± SEM and statistical analysis was done using two-way ANOVA **(B-F)**, ns *P>*0.05, \**P*<0.05, \*\**P*<0.01, \*\*\**P*<0.001, \*\*\*\**P*<0.0001.

### GSDMD-deficient platelets ameliorate excessive pulmonary thrombosis by restoring monocyte efferocytosis in influenza virus with secondary bacterial infection

To evaluate the relationship between platelet pyroptosis-driven procoagulant platelet and monocyte efferocytosis to the exacerbation of pulmonary thrombosis in severe pneumonia in *vivo*, we generated platelet-specific *Gsdmd*-deficient (*Gsdmd^flox/flox^* PF4-Cre) mice and subjected them, along with *Gsdmd^flox/flox^* littermate controls, to influenza virus and secondary bacterial challenge **(Figure 5A)**. Following infection, *Gsdmd^flox/flox^* PF4-Cre mice exhibited significantly reduced lung index, bacterial loads and pathological injury (including hyaline thrombi and inflammatory infiltration, black and red arrows in **Figure 5B**) compared with controls **(Figures 5B and 5C)**. BALF concentrations of TAT, sCD62P, IL-1β, IL-6 and TNF-α were substantially lower in the platelet-specific *Gsdmd* knockout group **(Figures 5D and S5)**. In addition, infected *Gsdmd^flox/flox^*PF4-Cre mice had higher circulating platelet counts, while pulmonary levels of Fibrin, TF and FXIIIa, as well as platelet-associated CD62P, CD40L, IL-1β and TNF-α in BALF, were markedly downregulated **(Figures 5E-5G)**. These results suggest that pyroptotic signaling mediated by platelet GSDMD exacerbates pulmonary thrombosis and lung injury in severe pneumonia following influenza with secondary bacterial infection. Further characterization of monocytes and efferocytosis receptors revealed that *Gsdmd^flox/flox^* PF4-Cre mice had higher proportions of F4/80^-^ CD11b^+^Ly-6C^+^, CX3CR1^+^Ly-6C^+^, and Mertk^+^Ly-6C^+^ monocyte populations in both BALF and lung tissues **(Figures 5H and 5I)**. Collectively, these data imply that platelet GSDMD-driven procoagulant platelet is associated with impaired monocyte efferocytosis and contributes to pulmonary thrombus exacerbation and lung injury in severe pneumonia.

**Figure 5.**
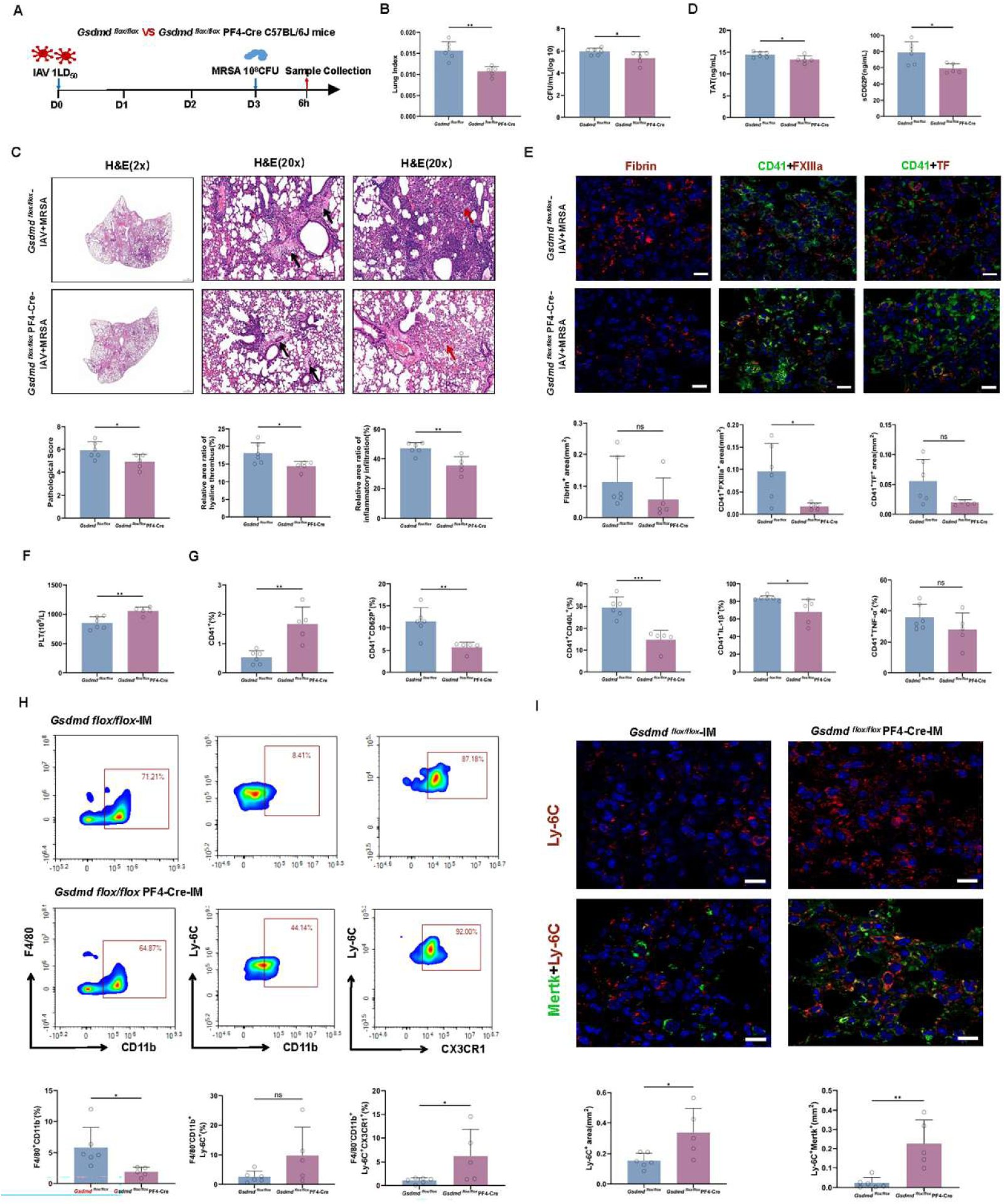
GSDMD-deficient platelets ameliorate excessive pulmonary thrombosis by restoring monocyte efferocytosis in influenza virus with secondary bacterial infection. **(A)** Schematic: Severe pneumonia of *Gsdmd^flox/flox^* mice *and Gsdmd^flox/flox^* PF4-Cre mice (Platelet specific knockdown of GSDMD mice) were induced by sequentially intranasal infection with IAV and MRSA (*Gsdmd^flox/flox^-* IAV+MRSA, n=6; *Gsdmd^flox/flox^* PF4-Cre*-*IAV+MRSA, n=5) to analyze the correlation of GSDMD-mediated procoagulant platelets with monocyte efferocytosis and pulmonary thrombus exacerbation. **(B)** The lung index and bacterial count in BALF of mice were calculated. **(C)** The transparent thrombus and inflammatory infiltration in lung tissue were detected by H&E staining, scale bars, 1000 or 100 μm. The black and red arrows indicate hyaline thrombi and inflammatory cell infiltration, respectively. **(D)** ELISA detection of thrombotic (TAT and sCD62P) factors expression in BALF. **(E)** Fibrin, FXIIIa and TF in mouse lung tissue were measured and semi quantitatively analyzed using immunofluorescence analysis and Image J calculation. Lung tissues were stained with nucleus for DAPI (blue), anti-Fibrin, FXIIIa and TF (red) and CD41 for platelets (green). Scale bars, 10 μm. **(F)** Blood counter detects the number of platelets in mouse blood. **(G)** FCM detects the number of platelets in BALF, as well as the expression of CD62P, CD40L, IL-1β and TNF-α derived from platelets. **(H and I) A significant decrease in efferocytosis receptor expression of monocytes was observed in *Gsdmd^flox/flox^* PF4-Cre mice after secondary infection with IAV and MRSA. (H)** BALF monocytes with efferocytosis receptor were analyzed via CD11b^+^F4/80^-^ Ly-6C^+^CX3CR1^+^ staining. **(I)** Lung cells co-expressing Ly-6C and Mertk were examined by IF staining. Data are presented as mean ± SEM and statistical analysis was done using Unpaired *t* test **(B-I)**, ns *P>*0.05, \**P*<0.05, \*\**P*<0.01.

### Procoagulant platelets attenuate monocyte efferocytosis by releasing complement C1qa

To identify factors secreted by procoagulant platelets that regulate monocyte efferocytosis, we comparatively analyzed the proteomic profiles of platelets stimulated with influenza virus alone versus those subjected to secondary bacterial challenge after influenza virus. REACTOME pathway enrichment analysis revealed that the "complement cascade" and "regulation of complement cascade" pathways were markedly enriched in platelets exposed to sequential infection compared with influenza virus alone. Among the pathway-associated proteins, complement components including C1qa, C1qb, C1qc, Masp2 and Mbl1 were significantly upregulated following dual stimulation **(Figure 6A)**. Given that platelet complement C1q and its subunits have been shown to regulate platelet activation and procoagulant activity^16, 17^, we focused our subsequent analysis on the complement C1q family, particularly C1qa. Our results showed that secondary bacterial challenge after influenza virus elevated complement C1qa expression in platelet lysates and supernatants in *vitro*, and also resulted in higher C1qa levels in BALF and enhanced pulmonary colocalization of C1qa with CD41 in *vivo*, compared with influenza virus alone **(Figures 6B, 6C, and S6A)**. These data indicate that secondary bacterial infection potently induces platelet C1qa expression and secretion.

**Figure 6.**
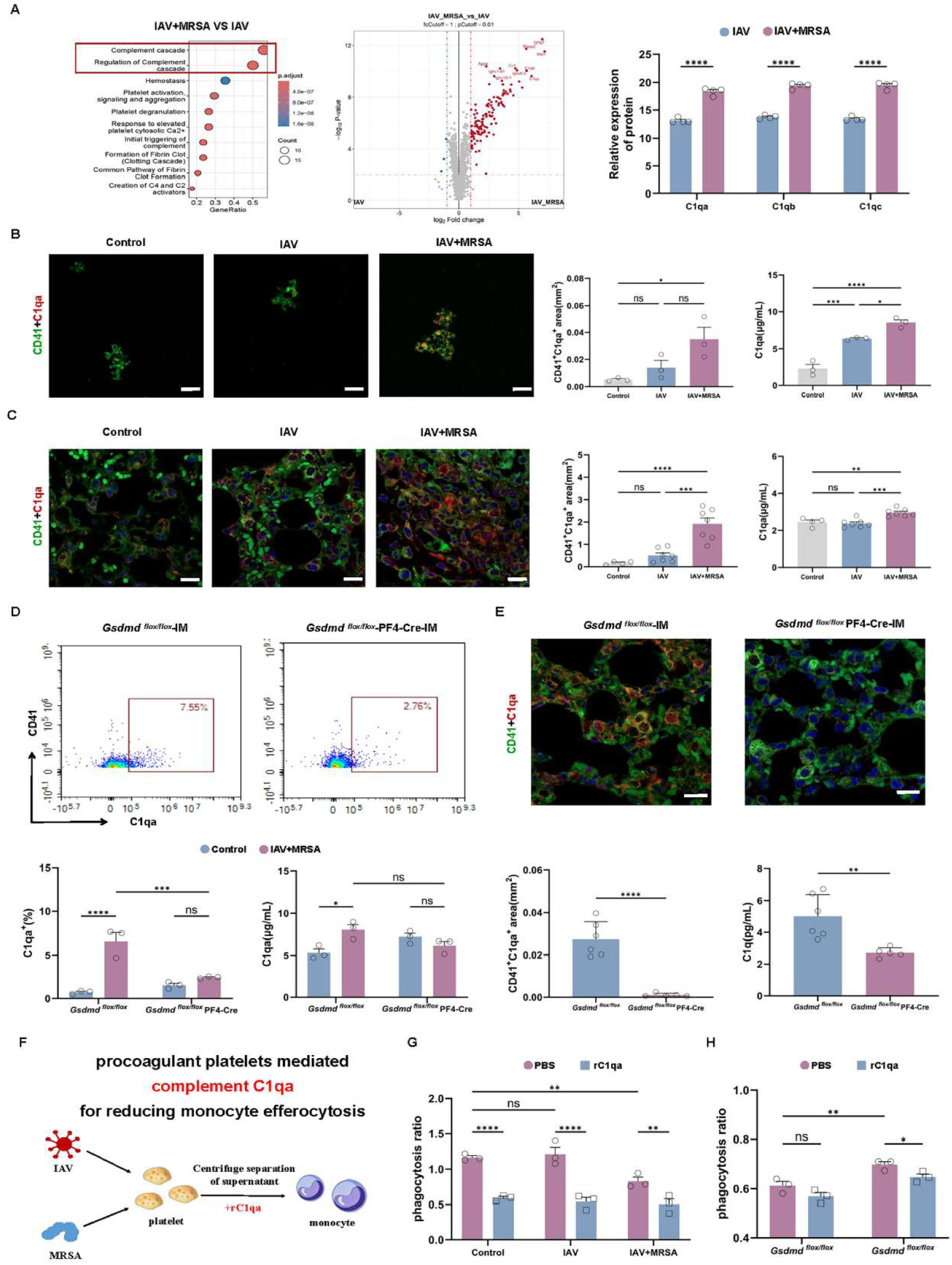
Procoagulant platelets attenuate monocyte efferocytosis by releasing complement C1qa. (A-**C) Increased production of platelet associated complement C1qa in severe pneumonia induced by influenza virus secondary bacterial infection. (A)** Over-Representation Analysis of Differential Proteins showed that “complement cascade” and “the regulation of complement cascade” pathways, as well as complement proteins such as C1qa, C1qb, C1qc, Masp2 and Mbl1, were significantly enriched, after platelet upon secondary stimulation with IAV and MRSA, compared to IAV stimulation (n=4). **(B)** Complement C1qa in the interior and supernatant of platelets was detected by immunofluorescence and ELISA (n=3. Scale bars, 10 μm). **(C)** The expression of platelet associated C1qa was detected in the lungs and BALF of mice with severe pneumonia following secondary challenge with IAV and MRSA (Control, n=4; IAV, n=7; IAV+MRSA, n=7. Scale bars, 10 μm). **(D-E) Influenza virus secondary bacterial infection induced platelet complement C1qa production dependent on necroptosis and pyroptosis. (D)** The detection of complement C1qa inside and supernatant of *Gsdmd^flox/flox^* and *Gsdmd^flox/flox^* PF4-Cre mice platelets after secondary stimulation with IAV and MRSA (n=3. Scale bars, 10 μm). **(E)** When *Gsdmd^flox/flox^* and *Gsdmd^flox/flox^*PF4-Cre mice were infected with IAV followed by secondary MRSA, detection of platelet associated C1qa in the lungs and C1qa in BALF was observed (*Gsdmd ^flox/flox^-*IAV+MRSA, n=6; *Gsdmd^flox/flox^* PF4-Cre*-*IAV+MRSA, n=5. Scale bars, 10 μm). **(F-H) The effect of complement C1qa on monocyte efferocytosis. (F)** In order to clarify whether the effect of soluble platelet supernatant on monocyte efferocytosis was dependent on complement C1qa, recombinant complement C1qa protein (rC1qa) also was added to the platelet supernatant after secondary stimulation with IAV and MRSA, and the efferocytic effect of monocyte was observed. **(G)** The reduced efferocytic activity of monocytes in platelet supernatant upon secondary stimulation with IAV and MRSA was further lessened by the addition of rC1qa (n=3). **(H)** Compared with the significantly reduced efferocytosis of platelet supernatant from *Gsdmd^flox/flox^* mice, the efferocytic fuction of *Gsdmd^flox/flox^* PF4-Cre mouse platelets on monocytes was restored after secondary stimulation with IAV and MRSA, while the efferocytic effect was reduced after adding rC1qa (n=3). Data are presented as mean ± SEM and statistical analysis was done using Unpaired *t* test **(A and E)** or one-way ANOVA **(B and C)** or two-way ANOVA **(D, G and H)**, ns *P>*0.05, \**P*<0.05, \*\**P*<0.01, \*\*\**P*<0.001, \*\*\*\**P*<0.0001.

To determine whether platelet necroptosis and pyroptosis contribute to C1qa upregulation, we performed pharmacological inhibition and genetic knockout experiments. Early administration of GSK-872 or NSA suppressed the elevation of complement C1qa induced by sequential infection, with NSA showing a more pronounced inhibitory effect **(Figure S6B)**. Platelets from *Gsdmd*^-/-^, *Nlrp3*^-/-^ and *Gsdmd^flox/flox^* PF4-Cre mice exhibited significantly reduced C1qa expression and secretion upon dual stimulation **(Figures 6D and S6C)**. Consistently, lower C1qa levels and reduced colocalization of C1qa with CD41 were observed in the BALF and lungs of *Gsdmd^flox/flox^* PF4-Cre mice **(Figure 6E)**. Collectively, these results demonstrate that complement C1qa production is highly dependent on NLRP3-GSDMD-mediated pyroptosis and RIPK3-MLKL-mediated necroptosis.

To directly establish that complement C1qa mediates the suppression of monocyte efferocytosis, we co-cultured monocytes with platelet supernatants derived from various stimulation conditions, with or without exogenous recombinant C1qa (rC1qa), and assessed efferocytic activity **(Figure 6F)**. Addition of rC1qa alone reduced monocyte efferocytosis to an extent comparable to that induced by supernatants from platelets subjected to influenza virus with secondary bacterial challenge. When rC1qa was further added to pyroptotic and necroptotic platelet supernatants, the suppression of efferocytosis was exacerbated **(Figure 6G)**. In rescue experiments, the restored efferocytosis observed in monocytes cultured with NSA-pretreated pyroptotic and necroptotic platelet supernatants was again compromised upon rC1qa supplementation **(Figure S6D)**. Similarly, the improved efferocytosis seen with supernatants from *Nlrp3*^-/-^ and *Gsdmd^flox/flox^*PF4-Cre mice platelets following sequential infection was markedly attenuated by rC1qa addition, ultimately recapitulating the impaired phenotype induced by wild-type or control supernatants **(Figures 6H and S6E)**. Collectively, these results demonstrate that supernatants from platelets undergoing necroptosis and pyroptosis suppress monocyte efferocytosis in a complement C1qa-dependent manner, and identify complement C1qa as a key mediator through which procoagulant platelets dampen efferocytosis.

### Procoagulant platelets and reduced efferocytosis receptor expression on monocytes in the BALF of severe pneumonia patients

To validate the clinical relevance of our findings and to determine whether procoagulant platelets and impaired monocyte efferocytosis are observed not only in influenza-associated secondary bacterial pneumonia but also in severe pneumonia of diverse viral or bacterial etiologies, we isolated platelets from patients with severe pneumonia and examined their morphology, procoagulant activity and programmed cell death pathways. TEM revealed that platelets from healthy controls exhibited uniform morphology, whereas those from patients with severe pneumonia displayed marked size heterogeneity, frequent swelling and necrosis **(Figure 7A)**. Platelets from these patients showed significantly upregulated expression of activated caspase-1 and cleaved GSDMD-N (a pyroptosis marker), as well as the p-RIPK1/p-RIPK3/p-MLKL signaling cascade (necroptosis markers) **(Figures 7B-7D)**. A significant increase in procoagulant platelets, defined as platelets with high expression of TF, CD62P and CD40L, were also observed. **(Figures 7D and 7E)**. Notably, platelets with high TF expression were significantly enriched in BALF from patients with severe pneumonia **(white arrows in Figure 7F)**, suggesting that necroptosis- and pyroptosis-mediated procoagulant activity occurs in severe pneumonia. In parallel, we characterized efferocytosis receptor expression on monocytes from these patients and examined the effects of patient-derived platelet supernatants on monocyte function. Monocytes with low CX3CR1 and Mertk expression were frequently detected in BALF **(yellow and orange arrows in Figure 7F)**. Compared with supernatants from healthy subjects, those from severe pneumonia patients significantly reduced efferocytosis, downregulated CX3CR1 expression and decreased IL-10 production in monocytes **(Figures S7)**. These data indicate that monocyte efferocytosis is impaired in severe pneumonia, and that supernatants from necrotic or swollen platelets derived from these patients directly attenuate efferocytosis. Collectively, these clinical data further corroborate the translational relevance of our experimental observations.

**Figure 7.**
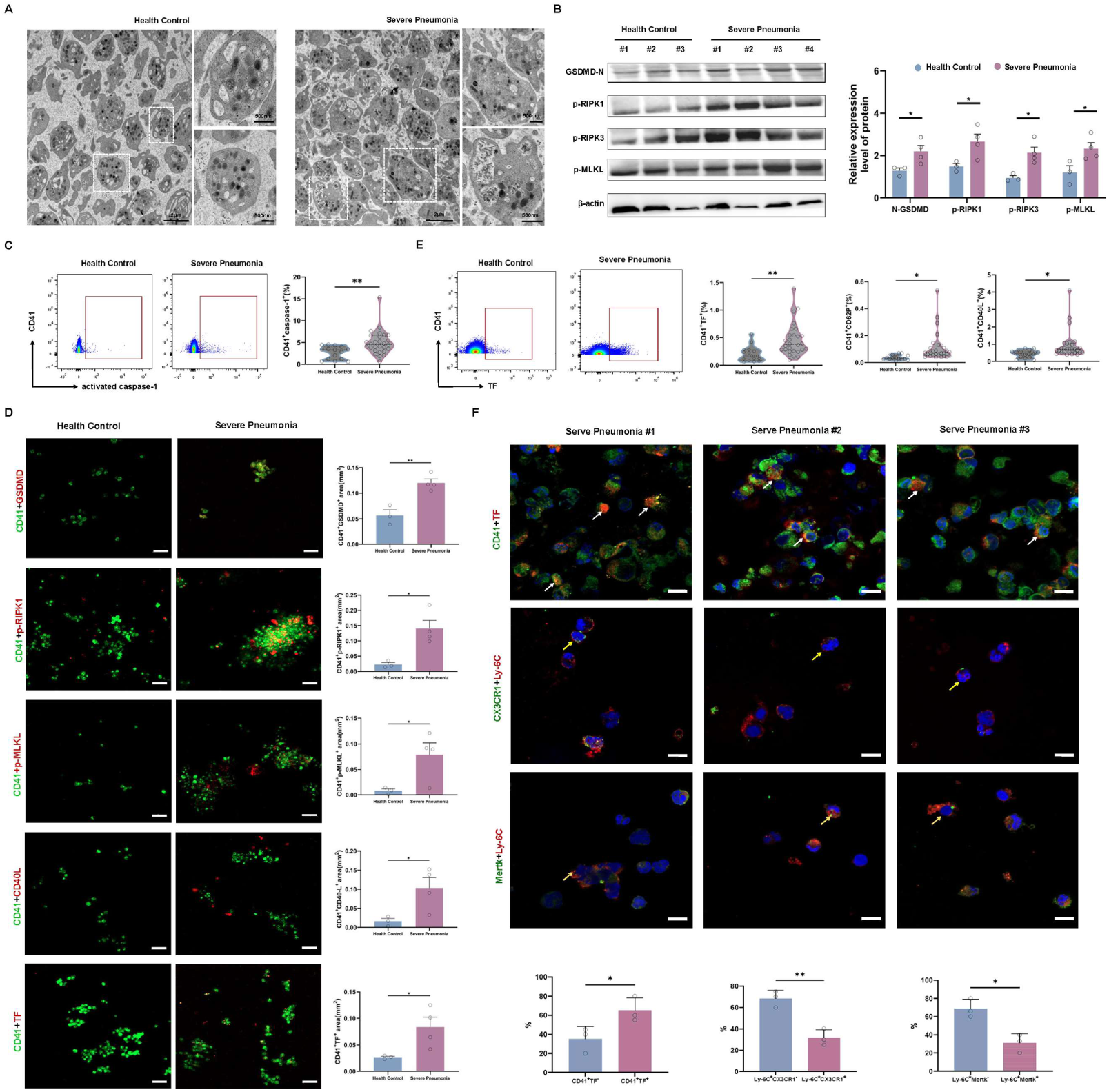
Procoagulant platelets and reduced efferocytosis receptor expression on monocytes in patients with severe pneumonia. **(A-E) Necrotic and pyroptosis platelets in the blood of patients with severe pneumonia. (A)** Transmission electron microscopy detection of platelet morphology in clinical healthy controls and severe pneumonia patients. Scale bars, 2 μm or 500 nm. **(B)** WB detection of GSDMD-N, p-RIPK1, p-RIPK3 and p-MLKL expression in platelets of patients with severe pneumonia (Health Control, n=3; Severe Pneumonia, n=4). **(C)** Flow cytometric analysis revealed significantly elevated levels of active caspase-1 in platelets derived from patients with severe pneumonia (Health Control, n=15; Severe Pneumonia, n=28). **(D)** Immunofluorescence detection of platelet GSDMD, p-RIPK1, p-MLKL, CD40L and TF expression in blood. Scale bars, 10 μm (Health Control, n=3; Severe Pneumonia, n=4). **(E)** Flow cytometry showed the expression of platelet-associated TF, CD62P and CD40L (Health Control, n=15; Severe Pneumonia, n=28). **(F) Procoagulant platelets and reduced efferocytosis receptor expression on monocytes in the BALF of patients with severe pneumonia.** Immunofluorescence detected platelet-associated TF expression and monocyte-associated CX3CR1 or Mertk expression in BALF of severe pneumonia patients (Severe Pneumonia, n=3). The white, yellow and orange arrows respectively represented TF-high platelets (CD41), CX3CR1-low monocytes (Ly-6C), and Mertk-low monocytes(Ly-6C). Scale bars, 10 μm. Data are presented as mean ± SEM and statistical analysis was done using Unpaired *t* test **(B-F)**, ns *P>*0.05, \**P*<0.05, \*\**P*<0.01.

## DISCUSSION

Using a mouse model of influenza A followed by MRSA, we demonstrate that MLKL-mediated necroptosis and GSDMD-mediated pyroptosis result in increased procoagulant platelets, which exacerbate pulmonary thrombosis and tissue damage through suppression of monocyte efferocytosis. We identify complement C1qa as a platelet-derived mediator linking these death pathways to impaired efferocytosis, and show that pharmacological inhibition or platelet-conditional *Gsdmd* knockout restores efferocytosis and attenuates thrombotic and pulmonary damage. Clinical relevance is supported by bronchoalveolar lavage fluid from severe pneumonia patients, in which procoagulant platelets from both death pathways and diminished efferocytic receptor levels on monocytes are detected. Based on these findings, we speculate that in influenza virus infection alone, platelets may undergo apoptosis and promote monocyte efferocytosis, thereby acting as a brake to restrain progressive thrombus exacerbation and resulting in only mild injury. In contrast, following secondary bacterial infection, platelets undergo necroptosis and pyroptosis, leading to increased procoagulant platelets that provide a catalytic surface for coagulation factor assembly and thrombin generation. Concomitantly, monocyte efferocytosis is impaired, and this brake is lost, leading to a progressive thrombotic cascade and aggravated injury **(Graphic Abstract)**. Collectively, this work identifies monocyte efferocytosis as a previously unrecognized mechanism by which procoagulant platelets aggravate thrombosis, extending platelet-monocyte crosstalk from thrombus initiation to thrombus exacerbation and positioning procoagulant platelets, monocyte efferocytosis and complement C1qa as a potential therapeutic strategy for thrombus-exacerbating diseases.

**Graphic Abstract.**
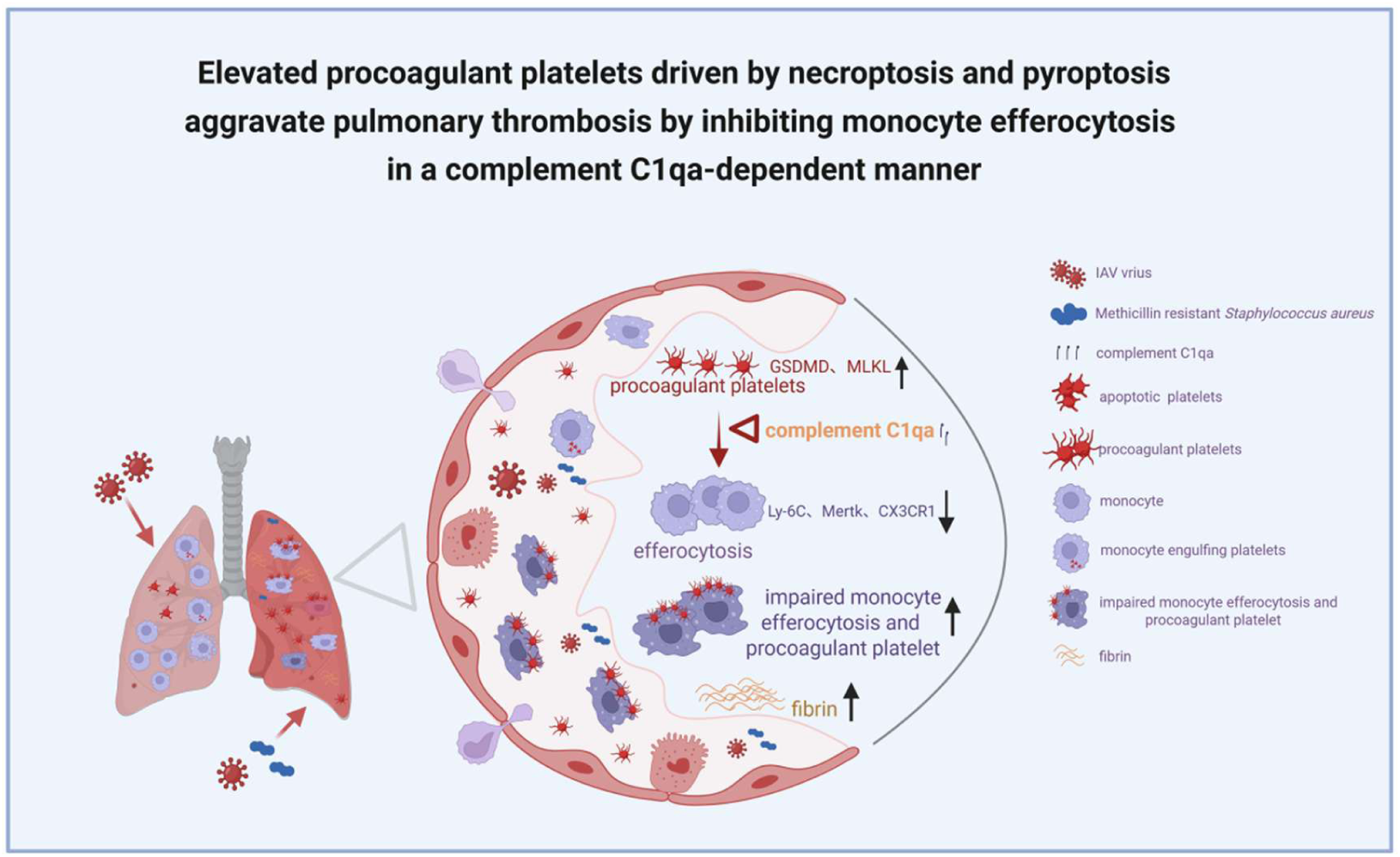
Elevated procoagulant platelets driven by necroptosis and pyroptosis aggravate pulmonary thrombosis by inhibiting monocyte efferocytosis in a complement C1qa-dependent manner during severe pneumonia. In influenza virus infection alone, platelets may undergo apoptosis and promote monocyte efferocytosis, thereby acting as a brake to restrain progressive thrombus exacerbation and resulting in only mild injury. In contrast, following secondary bacterial infection, platelets undergo necroptosis and pyroptosis, leading to increased procoagulant platelets that provide a catalytic surface for coagulation factor assembly and thrombin generation. Concomitantly, monocyte efferocytosis is impaired, and this brake is lost, leading to a progressive thrombotic cascade and aggravated injury.

Pulmonary thrombus exacerbation is a key pathological feature of severe pneumonia, particularly in influenza virus pneumonia with secondary bacterial infection, and is a major contributor to disease progression and mortality^2^. Procoagulant platelets, a subpopulation characterized by surface phosphatidylserine (PS) and P-selectin expression, are increasingly recognized as key effectors in thrombotic diseases and as contributors to pulmonary thrombus exacerbation^7^. Their prothrombotic actions are mediated primarily through PS externalization, which provides a catalytic surface for tenase and prothrombinase complex assembly and enhances thrombin generation and clot formation^18^. Additionally, procoagulant platelets release PS-bearing microvesicles that amplify coagulation signaling^19^. Platelet-monocyte crosstalk has been implicated in thrombus initiation, with P-selectin and CD40L driving monocyte tissue factor expression and procoagulant activity^8, 9^. However, the mechanisms underlying procoagulant platelet generation in specific pathological contexts, and how their interaction with monocytes transitions from thrombus initiation to thrombus exacerbation, remain incompletely understood.

Platelet programmed cell death pathways are emerging as potential sources of procoagulant activity, and accumulating evidence suggests that distinct death modalities (apoptosis, necroptosis, and pyroptosis) may differentially contribute to thrombus formation and propagation^20^. Prior studies in other infections have shown that different pathogens activate distinct platelet death pathways: severe fever with thrombocytopenia syndrome virus triggers pyroptosis, apoptosis, necroptosis and autophagy^21^; dengue virus induces platelet activation with elevated CD62P and NLRP3-caspase-1-IL-1β signaling^22^; and *E. coli* α-hemolysin elicits a necrotic and procoagulant phenotype^23^. However, whether and which death pathways generate procoagulant platelets in severe pneumonia following influenza virus with secondary bacterial infection had not been defined. In our model, influenza virus alone induced platelet shrinkage, whereas secondary bacterial infection triggered both MLKL-mediated necroptosis and GSDMD-mediated pyroptosis, leading to platelet swelling, rupture, and procoagulant phenotype. Importantly, platelet-conditional *Gsdmd* deficiency markedly reduced procoagulant platelet formation and correlated with reduced thrombotic burden, indicating that these two programmed death pathways are critical for generating the procoagulant platelet population, thereby exacerbating pulmonary thrombosis in influenza-associated secondary bacterial pneumonia.

Platelets undergoing distinct death modes engage immune cells to modulate inflammation and thrombosis, thereby maintaining host homeostasis or driving disease progression. For example, dengue-induced apoptotic platelets are efficiently cleared by macrophages via PS recognition^24^, and apoptotic platelets expressing FasL limit secondary inflammation in stroke models^25^. Collagen-stimulated activated platelets secrete prostaglandin E2, promoting IL-10 and reducing TNF-α production in monocytes^26^. These observations suggest that contracted or apoptotic platelets tend to maintain systemic homeostasis. In contrast, pyroptotic or necroptotic platelets seem to drive pathological inflammatory and prothrombotic cascades: GSDMD-mediated pyroptotic platelets facilitate NET formation and S100A8/A9 release in sepsis^15^, while cyclophilin D-dependent necrotic platelets provide a procoagulant surface for Fibrin deposition and promote neutrophil macroaggregates in gut ischemia^27, 28^. However, whether these platelet-driven pathways contribute to the subsequent growth of established thrombi, particularly through modulation of monocyte efferocytosis, remains largely unexplored. Monocyte polarization is known to influence thrombus fate, inflammatory subsets (CCR2^+^Ly-6C^+^) promote formation^29^, whereas pro-resolving subsets (Ly-6C^-^) facilitate resolution^30^. Yet the specific mechanisms governing thrombus exacerbation beyond the initial trigger, particularly those involving procoagulant platelets and monocyte efferocytosis, are incompletely defined. Our findings indicate that GSDMD-mediated pyroptotic platelets and MLKL-dependent necrotic platelets, via C1qa, dampen efferocytosis and also reduce the proportion of reparative (M2-like) monocytes, thereby associating with thrombus exacerbation.

The reparative monocytes are characterized by high efferocytic potential and are known to derive from inflammatory Ly-6C^+^ monocytes during atherosclerosis regression^31, 32^. This observation suggests that procoagulant platelets may impair efferocytosis through both a direct functional inhibition and a reduction in the cellular pool competent for clearance, thus associating with thrombus exacerbation. Such a dual mechanism is distinct from their classical role in clot initiation and highlights a new layer of platelet-monocyte crosstalk in thrombus exacerbation.

To identify the factors derived from procoagulant platelets that are responsible for the suppression of monocyte efferocytosis, we performed unbiased proteomic analysis and identified complement C1qa as a major mediator produced predominantly by necroptotic and pyroptotic platelets. While C1q subunits associated with platelets have been proposed as diagnostic biomarkers for coagulation disorders including sepsis, deep vein thrombosis and immune thrombocytopenia^33-35^, and C1q can modulate platelet CD62P expression and promote procoagulant activity^16, 17^, its cellular source and functional impact on monocyte efferocytosis in pneumonia have not been previously addressed. Our study fills this gap by showing that platelet C1qa production is tightly coupled to necroptotic and pyroptotic pathways, extending to the platelet compartment the known connection between programmed cell death and complement activation previously described for fibroblast necroptosis and for complement generation mediated by NETs^36, 37^. Importantly, we observe that excess C1qa impairs monocyte efferocytosis in *vitro*, and this effect is abrogated when platelets lacking *Gsdmd* are used. This finding appears to contrast with earlier reports that complement C1q binding to apoptotic cells enhances efferocytosis by monocytes^38^. However, those studies employed etoposide-induced apoptotic Jurkat cells, whereas our system involves necroptotic or pyroptotic platelets induced by pathogen challenge. The discrepancy likely reflects differences in C1qa function that depend on cell type and stimulus, and may also suggest that C1qa derived from distinct cellular sources exerts opposing effects. In support of our results, single-cell RNA-sequencing analysis in a murine pneumonia model revealed that high C1qa expression in monocyte-derived macrophages correlates with impaired bacterial clearance^39^, although the role of platelets was not examined in that work. Collectively, our findings establish a previously unrecognized link between platelet death pathways, complement C1qa and monocyte efferocytosis, deepening the understanding of complement-mediated regulation in infectious thrombosis.

The presence of procoagulant platelets (derived from both MLKL- and GSDMD-dependent pathways) and reduced CX3CR1 and Mertk expression on monocytes in BALF from patients with severe pneumonia corroborates our murine findings. Platelet supernatants from patients with severe pneumonia also directly suppressed monocyte efferocytosis receptor levels, directly demonstrating the interaction between procoagulant platelets and monocyte efferocytosis. Given that apoptotic platelets often limit inflammation, whereas necroptotic/pyroptotic platelets drive pathological cascades, our study highlights a distinct prothrombotic mechanism that promotes thrombus exacerbation, centred on efferocytosis suppression. Interventions that inhibit procoagulant platelet formation or restore efferocytic capacity may offer therapeutic strategies, though further validation is needed. Overall, this work shifts focus from thrombus initiation to progression, positioning procoagulant platelets, monocyte efferocytosis and complement C1qa as key determinants in immunothrombosis of severe pneumonia and sepsis.

## Data and code availability

This paper does not report original code. Any additional information required to reanalyze the data reported in this paper is available from the lead contact upon request.

## Supporting information

This article contains supporting information.

## Acknowledgements

Thank you for the mice of *Nlrp3*^-/-^, *Gsdmd*^-/-^ and platelet-specific *Gsdmd* knockout gifted by Jing-Lin Wang and Yuan Yuan’s laboratory. We thank every member of Dr Zi-Feng Yang’s, Xiao-Hong Chen’s and Shu-Feng Ma’s lab for their helpful discussion and advice. This work was funded by Noncommunicable Chronic Diseases-National Science and Technology Major Project (2024ZD0528801 to Z.F.Y. and X.H.C.), National Multidisciplinary Innovation Team Project of Traditional Chinese Medicine (ZYYCXTD-D-202406 to Z.F.Y.), Yangcheng Traditional Chinese Medicine Innovative Talent Team Project (2026RC010 to Z.F.Y.), Special Support Program for Outstanding Talents of Guangdong Province (2024JC08Y008 to Z.F.Y.) and the grant of State Key Laboratory of Respiratory Disease (SKLRD-Z-202502 to S.F.M.).

## Author contributions

Conceptualization: Z.F.Y., Z.H.Z., S.F.M., X.H.C., Z.S.L. and S.Q.L.; Methodology: S.Q.L., W.Y.F., J.B.L., X.X.L., M.Y.L., C.Y.K., H.X.Z. and L.C.Y.; Validation: S.Q.L., Y.Y.L., W.Y.F. and J.H.H.; Formal analysis: S.Q.L. and W.Y.F.; Investigation: Z.F.Y., Z.H.Z., S.F.M., X.H.C. and S.Q.L.; Data curation: S.Q.L., W.Y.F. and J.B.L.; Resources: Y.Y.L. and J.H.H.; Funding acquisition: Z.F.Y., S.F.M., X.H.C.; Writing—original draft preparation: S.Q.L.; Writing review and editing: S.Q.L., W.Y.F., Z.S.L., X.H.C. and S.F.M. All authors have read and agreed to the published version of the manuscript. The authors declare no conflict of interest.

## References

1. Taubenberger JK, Kash JC and Morens DM. The 1918 influenza pandemic: 100 years of questions answered and unanswered. Sci Transl Med. 2019;11:eaau5485.doi:10.1126/scitranslmed.aau5485

2. Sheng Z-M, Chertow DS, Ambroggio X, McCall S, Przygodzki RM, Cunningham RE, Maximova OA, Kash JC, Morens DM and Taubenberger JK. Autopsy series of 68 cases dying before and during the 1918 influenza pandemic peak. Proc Natl Acad Sci U S A. 2011;108:16416–21.doi:10.1073/pnas.1111179108

3. Walters K-A, D’Agnillo F, Sheng Z-M, Kindrachuk J, Schwartzman LM, Kuestner RE, Chertow DS, Golding BT, Taubenberger JK and Kash JC. 1918 pandemic influenza virus and Streptococcus pneumoniae co-infection results in activation of coagulation and widespread pulmonary thrombosis in mice and humans. J Pathol. 2016;238:85–97.doi:10.1002/path.4638

4. Yang Y and Tang H. Aberrant coagulation causes a hyper-inflammatory response in severe influenza pneumonia. Cellular & molecular immunology. 2016;13:432–42.doi:10.1038/cmi.2016.1

5. Yan M, Wang Z, Qiu Z, Cui Y and Xiang Q. Platelet signaling in immune landscape: comprehensive mechanism and clinical therapy. Biomarker research. 2024;12:164.doi:10.1186/s40364-024-00700-y

6. Tian Y, Zong Y, Pang Y, Zheng Z, Ma Y, Zhang C and Gao J. Platelets and diseases: signal transduction and advances in targeted therapy. Signal transduction and targeted therapy. 2025;10:159.doi:10.1038/s41392-025-02198-8

7. Denorme F and Campbell RA. Procoagulant platelets: novel players in thromboinflammation. Am J Physiol Cell Physiol. 2022;323:C951–c958.doi:10.1152/ajpcell.00252.2022

8. Eugenio D H, Isaclaudia G A-Q, Lohanna P, Lívia T, Ester A B, Camila R R P, Cassia R, Sérgio F, Thiago M L S, Pedro K, Fernando A B and Patrícia T BJB. Platelet activation and platelet-monocyte aggregate formation trigger tissue factor expression in patients with severe COVID-19. 2020;136.doi:10.1182/blood.2020007252

9. Li T, Yang Y, Li Y, Wang Z, Ma F, Luo R, Xu X, Zhou G, Wang J, Niu J, Lv G, Crispe IN and Tu Z. Platelets mediate inflammatory monocyte activation by SARS-CoV-2 spike protein. The Journal of clinical investigation. 2022;132.doi:10.1172/jci150101

10. Li W, Huang Y, Liu J, Zhou Y, Sun H, Fan Y and Liu F. Defective macrophage efferocytosis in advanced atherosclerotic plaque and mitochondrial therapy. Life Sci. 2024;359:123204.doi:10.1016/j.lfs.2024.123204

11. S R L, S M A, S L G, S M T, M G P, T D, S M L, P M H and C V JJCDD. Ly6C(+) monocyte efferocytosis and cross-presentation of cell-associated antigens. 2016;23.doi:10.1038/cdd.2016.24

12. Kyrmizi I, Ioannou M, Hatziapostolou M, Tsichlis PN, Boumpas DT and Tassiulas I. Tpl2 kinase regulates FcγR signaling and immune thrombocytopenia in mice. Journal of leukocyte biology. 2013;94:751–7.doi:10.1189/jlb.0113039

13. Sharma L, Wu W, Dholakiya SL, Gorasiya S, Wu J, Sitapara R, Patel V, Wang M, Zur M, Reddy S, Siegelaub N, Bamba K, Barile FA and Mantell LL. Assessment of phagocytic activity of cultured macrophages using fluorescence microscopy and flow cytometry. Methods in molecular biology (Clifton, NJ). 2014;1172:137–45.doi:10.1007/978-1-4939-0928-5_12

14. Mohammad E, Paresh P K, Vipin S, Susheel N C, Saroj Kant M, Rameshwar Nath C and Debabrata DJCDD. Necroptosis executioner MLKL plays pivotal roles in agonist-induced platelet prothrombotic responses and lytic cell death in a temporal order. 2023;30.doi:10.1038/s41418-023-01181-6

15. Meiling S, Chaofei C, Shaoying L, Musheng L, Zhi Z, Yuan Z, Luoxing X, Xiuzhen L, Dezhong Z, Qiqi L, Xuejiao F, Ying W, Yingying L, Feiyan C, Wei L, Yun B, Jinhong Q, Manli G, Miaoyun Q, Lei S, Renjing L, Ping W, John H and Wai Ho TJNCR. Gasdermin D-dependent platelet pyroptosis exacerbates NET formation and inflammation in severe sepsis. 2022;1.doi:10.1038/s44161-022-00108-7

16. Skoglund C, Wetterö J, Tengvall P and Bengtsson T. C1q induces a rapid up-regulation of P-selectin and modulates collagen- and collagen-related peptide-triggered activation in human platelets. Immunobiology. 2010;215:987–995.doi:10.1016/j.imbio.2009.11.004

17. Peerschke EI, Reid KB and Ghebrehiwet B. Platelet activation by C1q results in the induction of alpha IIb/beta 3 integrins (GPIIb-IIIa) and the expression of P-selectin and procoagulant activity. The Journal of experimental medicine. 1993;178:579–87.doi:10.1084/jem.178.2.579

18. Veuthey L, Aliotta A, Calderara D, Portela C and Alberio LJIJMS. Mechanisms Underlying Dichotomous Procoagulant COAT Platelet Generation-A Conceptual Review Summarizing Current Knowledge. 2022;23:2536.doi:10.3390/ijms23052536

19. Fiore M, Guy A and James CJJVE. Procoagulant Platelet Characterization by Measuring Phosphatidylserine Exposure and Microvesicle Release from Human Purified Platelets. 2024:39671342.doi:10.3791/67042

20. Li C, Braun A, Zu J, Gudermann T, Mammadova-Bach E and Anders HJ. Converging Molecular Mechanisms of Nucleated Cell Death Pathways and Procoagulant Platelet Formation. Cells. 2025;14.doi:10.3390/cells14141075

21. Fang Y, Shen S, Zhang J, Xu L, Wang T, Fan L, Zhu Q, Xiao J, Wu X, Jin J, Wu Q, Zhang Y, Tang S, Zheng X and Deng F. Thrombocytopenia in Severe Fever with Thrombocytopenia Syndrome Due to Platelets With Altered Function Undergoing Cell Death Pathways. The Journal of infectious diseases. 2025;231:e183–e194.doi:10.1093/infdis/jiae355

22. Hottz ED, Lopes JF, Freitas C, Valls-de-Souza R, Oliveira MF, Bozza MT, Da Poian AT, Weyrich AS, Zimmerman GA, Bozza FA and Bozza PT. Platelets mediate increased endothelium permeability in dengue through NLRP3-inflammasome activation. Blood. 2013;122:3405–14.doi:10.1182/blood-2013-05-504449

23. Pérez Vázquez K, Tau J, Leal Denis MF, Fader CM, Ostuni MA, Schwarzbaum PJ and Herlax V. Alpha hemolysin of Escherichia coli induces a necrotic-like procoagulant state in platelets. Biochimie. 2024;227:1–14.doi:10.1016/j.biochi.2024.06.001

24. Alonzo MT, Lacuesta TL, Dimaano EM, Kurosu T, Suarez LA, Mapua CA, Akeda Y, Matias RR, Kuter DJ, Nagata S, Natividad FF and Oishi K. Platelet apoptosis and apoptotic platelet clearance by macrophages in secondary dengue virus infections. The Journal of infectious diseases. 2012;205:1321–9.doi:10.1093/infdis/jis180

25. Schleicher RI, Reichenbach F, Kraft P, Kumar A, Lescan M, Todt F, Göbel K, Hilgendorf I, Geisler T, Bauer A, Olbrich M, Schaller M, Wesselborg S, O’Reilly L, Meuth SG, Schulze-Osthoff K, Gawaz M, Li X, Kleinschnitz C, Edlich F and Langer HF. Platelets induce apoptosis via membrane-bound FasL. Blood. 2015;126:1483–93.doi:10.1182/blood-2013-12-544445

26. Linke B, Schreiber Y, Picard-Willems B, Slattery P, Nüsing RM, Harder S, Geisslinger G and Scholich K. Activated Platelets Induce an Anti-Inflammatory Response of Monocytes/Macrophages through Cross-Regulation of PGE(2) and Cytokines. Mediators of inflammation. 2017;2017:1463216.doi:10.1155/2017/1463216

27. Hua VM, Abeynaike L, Glaros E, Campbell H, Pasalic L, Hogg PJ and Chen VM. Necrotic platelets provide a procoagulant surface during thrombosis. Blood. 2015;126:2852–62.doi:10.1182/blood-2015-08-663005

28. Yuan Y, Alwis I, Wu MCL, Kaplan Z, Ashworth K, Bark D, Pham A, Mcfadyen J, Schoenwaelder SM, Josefsson EC, Kile BT and Jackson SPJSTM. Neutrophil macroaggregates promote widespread pulmonary thrombosis after gut ischemia. 2017;9:eaam5861.doi:10.1126/scitranslmed.aam5861

29. Shahneh F, Christian Probst H, Wiesmann SC, N AG, Ruf W, Steinbrink K, Raker VK and Becker C. Inflammatory Monocyte Counts Determine Venous Blood Clot Formation and Resolution. Arterioscler Thromb Vasc Biol. 2022;42:145–155.doi:10.1161/atvbaha.121.317176

30. Kimball AS, Obi AT, Luke CE, Dowling AR, Cai Q, Adili R, Jankowski H, Schaller M, Holinstadt M, Jaffer FA, Kunkel SL, Gallagher KA and Henke PK. Ly6CLo Monocyte/Macrophages are Essential for Thrombus Resolution in a Murine Model of Venous Thrombosis. Thromb Haemost. 2020;120:289–299.doi:10.1055/s-0039-3400959

31. Rodríguez-Morales P and Franklin RA. Macrophage phenotypes and functions: resolving inflammation and restoring homeostasis. Trends Immunol. 2023;44:986–998.doi:10.1016/j.it.2023.10.004

32. Rahman K, Vengrenyuk Y, Ramsey SA, Vila NR, Girgis NM, Liu J, Gusarova V, Gromada J, Weinstock A, Moore KJ, Loke P and Fisher EA. Inflammatory Ly6Chi monocytes and their conversion to M2 macrophages drive atherosclerosis regression. J Clin Invest. 2017;127:2904–2915.doi:10.1172/jci75005

33. Zhang Y, Wang L, Kuang X, Tang D and Zhang P. Diagnostic and Prognostic Value of C1q in Sepsis-Induced Coagulopathy. Clinical and Applied Thrombosis/Hemostasis. 2024;30.doi:10.1177/10760296241257517

34. Xu Y, Zhang H, Jiao X, Zhang Y, Yin G, Wang C, Du Z, Liang M, Gao X, Gu Z, Jiang Y, Du B and Bi X. Dysregulations of C1QA, C1QB, C1QC and C5AR1 as candidate biomarkers of vascular dementia. npj Aging. 2025;11.doi:10.1038/s41514-025-00228-x

35. Lood C, Eriksson S, Gullstrand B, Jönsen A, Sturfelt G, Truedsson L and Bengtsson AA. Increased C1q, C4 and C3 deposition on platelets in patients with systemic lupus erythematosus – a possible link to venous thrombosis? Lupus. 2012;21:1423-1432.doi:10.1177/0961203312457210

36. Lusthaus M, Mazkereth N, Donin N and Fishelson Z. Receptor-Interacting Protein Kinases 1 and 3, and Mixed Lineage Kinase Domain-Like Protein Are Activated by Sublytic Complement and Participate in Complement-Dependent Cytotoxicity. Frontiers in immunology. 2018;9.doi:10.3389/fimmu.2018.00306

37. Schreiber A, Rousselle A, Becker JU, von Mässenhausen A, Linkermann A and Kettritz R. Necroptosis controls NET generation and mediates complement activation, endothelial damage, and autoimmune vasculitis. Proceedings of the National Academy of Sciences. 2017;114.doi:10.1073/pnas.1708247114

38. Fraser DA, Laust AK, Nelson EL and Tenner AJ. C1q Differentially Modulates Phagocytosis and Cytokine Responses during Ingestion of Apoptotic Cells by Human Monocytes, Macrophages, and Dendritic Cells. The Journal of Immunology. 2009;183:6175–6185.doi:10.4049/jimmunol.0902232

39. Stoner SN, Larson ED, Fulte S, Shaw SC, Fish ER, Janoff EN, Mack M and Clark SEJb. Myeloid cell reprogramming drives enhanced defense against Streptococcus pneumoniae lung infection following exposure to commensal Prevotella. 2026:2026.06.25.734552.doi:10.64898/2026.06.25.734552

